# SP1 amplifies the effects of cellular proliferation via the Wnt/β-catenin pathway in colon development and cancer

**DOI:** 10.1101/2025.06.05.658012

**Authors:** Ankita Sharma, Greg Jude Dsilva, Saurabh J. Pradhan, Sudiksha Mishra, Kirankumar Santhakumar, Sanjeev Galande

**Affiliations:** Department of Biology, Indian Institute of Science Education and Research, Dr Homi Bhabha Road, Pune 411008, India; Center of Excellence in Epigenetics, Department of Life Sciences, Shiv Nadar Institution of Eminence, Gautam Buddha Nagar, Uttar Pradesh 201314, India; Department of Genetic Engineering, SRM Institute of Science and Technology, Kattankulathur 603203, India

**Keywords:** SP1, β-catenin, Wnt signaling, cell-proliferation, colorectal cancer, zebrafish development

## Abstract

Wnt/β-catenin signaling is a highly conserved pathway across multicellular organisms, playing a pivotal role in various cellular processes essential for development and tissue homeostasis. Aberrant activation of this pathway is a major driver of colorectal cancer (CRC). The transcription factor SP1 has been identified as a key component of the Wnt/β-catenin signaling pathway through its physical interaction with β-catenin in colorectal cancer. However, the distinct roles played by this complex during development versus in disease contexts remain poorly understood. In this study, we conducted molecular and functional analyses by individually depleting SP1 and β-catenin. Interestingly, genes co-regulated by the SP1:β-catenin complex were significantly enriched for functions related to the cell cycle and DNA replication—key processes required for sustaining proliferation, whose disruption can promote tumorigenesis. These findings were further supported by data from mouse tumor models and clinical datasets. Additionally, cross-species analysis using zebrafish larvae revealed a synergistic interaction between SP1 and β-catenin, leading to excessive cell proliferation and developmental defects. Collectively, our results suggest a conserved role for the SP1:β-catenin complex in regulating cell proliferation during development and maintaining cellular homeostasis, processes critical for preventing cancer progression.

## Introduction

The Wnt/β-catenin signaling pathway is documented to regulate core cellular processes such as proliferation, differentiation, migration, and stem cell renewal^1^. It plays a crucial role in cell fate determination and axis formation during development, and its dysregulation is strongly implicated in disease, particularly colorectal cancer (CRC), where mutations in the Adenomatous Polyposis Coli (*APC*) gene account for over 70% of cases^2^. In the absence of Wnt ligands, β-catenin is targeted for degradation by the destruction complex, consisting of scaffold proteins APC and Axin; and kinases GSK3β and CK1α. Upon phosphorylation β-catenin is ubiquitinated by βTrCP and directed to proteasomal degradation^3^. On the contrary, when Wnt is activated, the ligand binds to the LRP5/6-Frizzled (Fzd) receptor complex, leading to the recruitment of Dishevelled (Dvl), which disrupts the destruction complex and stabilizesβ-catenin. Stabilized β-catenin translocates into the nucleus and interacts with TCF/LEF transcription factors to activate gene expression^1^.

While TCF/LEF proteins are primary partners for β-catenin, it also interacts with other transcriptional regulators including transcription factors (LRH-1, AR), chromatin modifiers (CBP, MLL1, Brg-1, ISW1) and the transcriptional co-activators (TBP, MED12, Parafibromin)^4^. Molecular functions of β-catenin occurring independently of TCF/LEF, results in the activation of distinct gene sets ^5,6^. Notably, members of the Specificity Protein (SP) family have emerged as important Wnt-associated transcription factors. Prior studies have shown that SP1 interacts and stabilizes β-catenin in CRC cell lines^7^. Upon Wnt activation, SP1 and β-catenin form a complex and translocate to the nucleus. In the absence of Wnt signaling, both proteins are degraded by the destruction complex. SP1:β-catenin complexes bind to Wnt target gene promoters, including *MYC* and *TCF7*^7^. Interestingly, this regulation occurs directly at the protein level and not at the transcriptional level (Figure 1A)^7^. In contrast, SP5 has been shown to act as a feedback inhibitor of Wnt signaling. Interestingly, Huggins et al. hypothesized that SP5 might be competing with SP1 for binding to a limited set of loci upon Wnt activation. These loci are mainly the Wnt target genes which are negatively regulated by SP5^8^. SP5 and SP8 have also been implicated in enhancer recruitment to β-catenin-LEF1 complexes at specific gene loci^9^.

**Figure 1:**
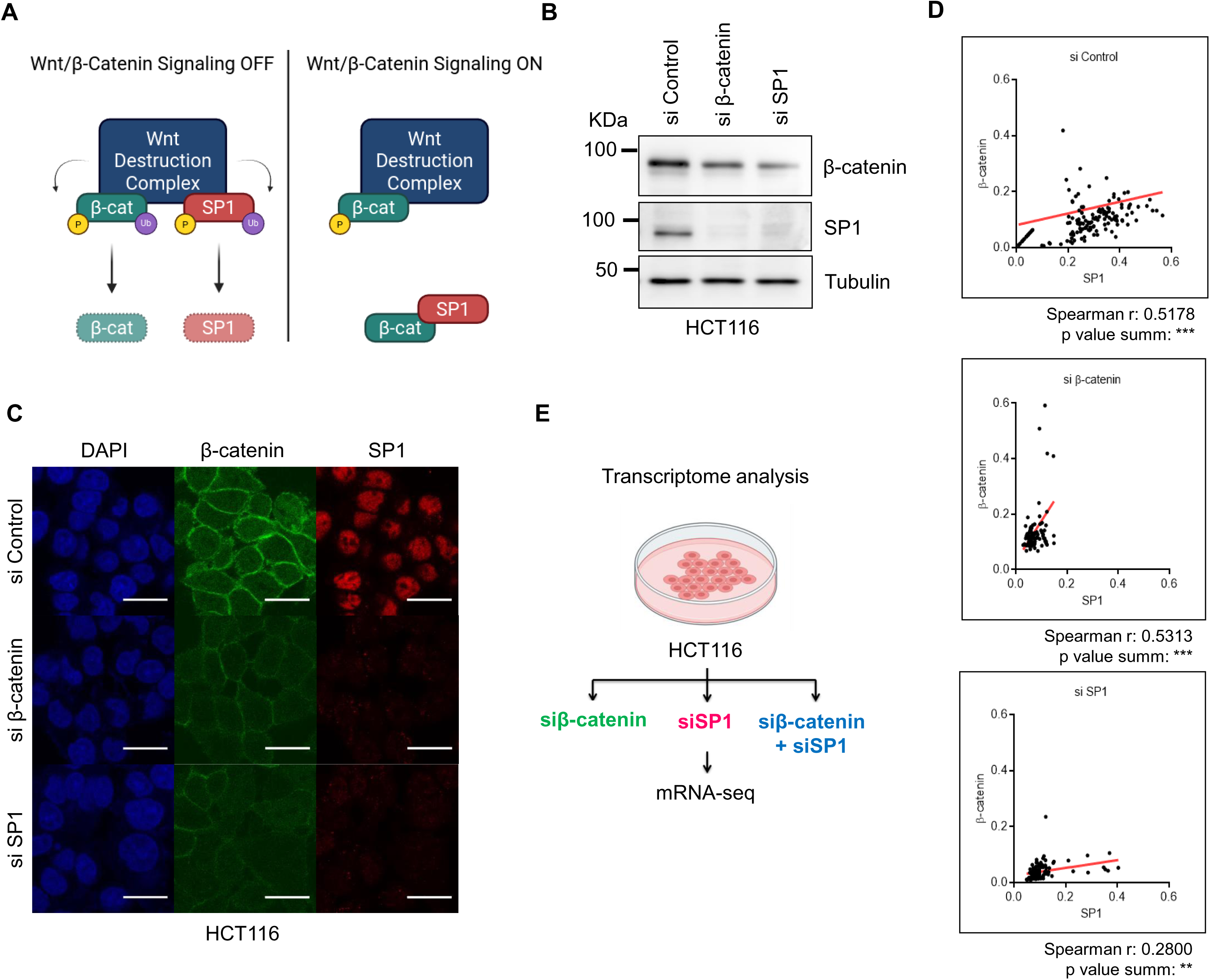
β-catenin and SP1 expression and perturbation show positive correlation in HCT116. **A.** Model for the regulation of SP1 via Wnt/β-catenin pathway. SP1 stability is regulated by the Wnt pathway. In the absence of Wnt activation, SP1 can get degraded via the activity of Wnt DC. Upon Wnt activation, SP1 gets stabilised along with the Wnt effector β-catenin. It also allows the interaction of SP1 and β-catenin which further aids in their stabilisation. **B.** Immunoblot for β-catenin and SP1 upon siβ-catenin and siSP1 treatment. **C.** Immunofluorescence image for β-catenin and SP1 upon siβ-catenin and siSP1 treatment. (Scale bar = 20 µM). **D.** Scatter plot for nuclear intensities for β-catenin and SP1 in cells treated with control siRNA (top), siβ-catenin (middle) and siSP1 (bottom). A regression line is depicted in red (Paired t-test performed to calculate significance). **E.** Schematic for the generation of transcriptome data.

To further investigate the SP1–β-catenin interaction, we performed transcriptomic profiling and chromatin occupancy analyses in HCT116 cells following individual and combined knockdowns of SP1 and β-catenin. Co-depletion led to a marked reduction in cell proliferation and increased apoptosis, with SP1 modulating both TCF-dependent and TCF-independent β-catenin target genes. In a mouse xenograft model, simultaneous inhibition of SP1 and β-catenin suppressed tumor growth more effectively than targeting either factor alone, underscoring their collaborative role in tumor progression. Chromatin occupancy analyses revealed that SP1 co-occupies regulatory regions alongside key components of the Wnt enhanceosome, contributing to its formation and activity. Parallel experiments in zebrafish demonstrated comparable synergistic effects during early development, emphasizing the evolutionary conservation of this regulatory mechanism. Collectively, these results highlight the SP1–β-catenin complex as a vital regulatory module of cell proliferation and survival in both developmental and oncogenic settings, particularly in colorectal cancer.

## Results

### Co-regulation of β-Catenin and SP1 in colorectal cancer cells

We previously showed that the depletion of either β-catenin or SP1 leads to the downregulation of the other protein, an epistasis phenomenon^7^. We validated these findings by using transient knockdown experiments and observed significant decrease in protein levels using immunoblotting (Figure 1B) and immunofluorescence assays (Figure 1C). Further, we estimated the degree of correlation between these two proteins by plotting nuclear staining intensities and observed positive correlation ensuring uniform depletion across the cells (Figure 1D). It is noteworthy that upon SP1 knockdown the correlation was weak as compared to other two conditions (control and siβ-catenin). This could be attributed to potential differences in the regulatory machinery getting affected in response to SP1 depletion as opposed to β-catenin depletion. Together, our results demonstrated that SP1 and β-catenin protein levels correlate with each other under both control as well as upon functional depletion.

### Synergistic transcriptional activity of β-Catenin and SP1

To understand how SP1–β-catenin complex regulate transcriptional changes in CRC cells, we performed transcriptome analysis upon siRNA-mediated depletion of β-catenin alone, SP1 alone, and β-catenin + SP1 (Schematic shown in Figure 1E). Efficiency of knockdown was confirmed at both RNA and protein levels (Supplementary Figure S1A&B). Since f all the siRNAs used in this study showed high efficiency, we selected siRNA #1 for β-catenin and siRNA #1 for SP1 (see methods for sequences). Additionally, we kept the amount of siRNA identical in all the conditions; therefore, the amount of each siRNA used in the case of double knockdown (DKD) was half of what was used in the case of single knockdown (SKD).

Upon β-catenin KD, although transcripts of β-catenin were significantly reduced, SP1 levels remained unchanged (Supplementary Figure 2A). Similarly, upon SP1 KD, β-catenin levels remained unaffected (Supplementary Figure S2F), confirming our previous findings that SP1 and β-catenin do not regulate each other transcriptionally but by inducing destabilization of each other^7^. RNA-seq analysis revealed 1304 upregulated and 1260 downregulated genes for β-catenin KD (Supplementary Figure S2B); and 2052 upregulated and 2071 downregulated genes upon SP1 KD (Supplementary Figure S2G). Further, β-catenin KD associated upregulated genes were enriched for lipid metabolism, metabolic processes, encompassing lipid catabolism, fatty acid and lipid metabolism, as well as lipid localization (Supplementary Figure S2C). It is worth noting that the Wnt/β-catenin pathway has been previously implicated in the direct activation of lipogenic genes in adipocytes^10,11^. β-catenin-activated genes that were downregulated were enriched for pathways involved in DNA replication, repair, and the cell cycle (Supplementary Figure S2D). Transcriptional regulatory relationship analysis (TRRUST) further revealed enrichment of both positive (e.g., SMAD3, MYC, JUN) and negative (e.g., E2F1, ESR1, TP53, SRY) regulators of Wnt/β-catenin signaling (Supplementary Figure S2E). SP1 targets were found among both upregulated and downregulated genes, suggesting its complex regulatory role. Collectively, our dataset confirmed known Wnt target gene regulation and offered a comprehensive view of β-catenin’s transcriptomic impact, identifying classical targets like *AXIN2* and *TCF7* alongside genes involved in metabolism and cell cycle control (Supplementary Figure S2O). Transcriptome analysis of SP1 KD HCT116 cells showed that upregulated genes were enriched for pathways involving TP63, TP53, cytochrome c release, and autophagy, suggesting SP1 loss induces apoptosis via TP53 (Supplementary Figure S2H). Downregulated genes were associated with the cell cycle, DNA replication, and repair (Supplementary Figure S2I). TRRUST analysis indicated enrichment of both pro-death (TP53, TP73) and pro-proliferation (E2F1, MYCN) regulators, highlighting SP1’s dual role in promoting proliferation while suppressing apoptosis (Supplementary Figure S2J). Key genes affected included DAPK3, CASP4, and CCNA2 (Supplementary Figure S2P).

Given the established roles of SP1 and Wnt/β-catenin in colorectal cancer stem cells (CCSCs)^12^, we further investigated their combined transcriptional activity in this context^12,13^. Combinatorial knockdown affected the cellular morphology and growth in a manner similar to SKD (Supplementary Figure S1C & Figure 2E). The quantification of both β-catenin and SP1 transcripts in this specific dataset revealed a decrease at the transcript level (Supplementary Figure S2K). In comparison to the SKD cells, the number of dysregulated genes was slightly higher, with 2683 genes displaying increased expression and 2275 genes exhibiting decreased expression in comparison to the control condition (Figure 2A). For further analysis, we overlapped the complete set of differentially expressed (DE) genes from each of the transcriptome datasets generated thus far (i.e., siβ-catenin, siSP1, siβ-catenin + siSP1) to identify a common set of genes (Figure 2B). This examination revealed that 1837 genes exhibited differential expression in all three datasets. Apart from these, 1591 genes were not picked up in any SKD datasets. We proceeded with the genes that were common in all three datasets. Next, we separately plotted the expression change values (in terms of log2 Fold Change) of the genes under the positive and negative regulation of β-catenin and SP1. We did observe that the level of increase or decrease in expression was significantly higher in the DKD condition as compared to any of the SKD conditions, pointing towards a synergistic effect in their transcriptional activities (Figure 2C).

**Figure 2:**
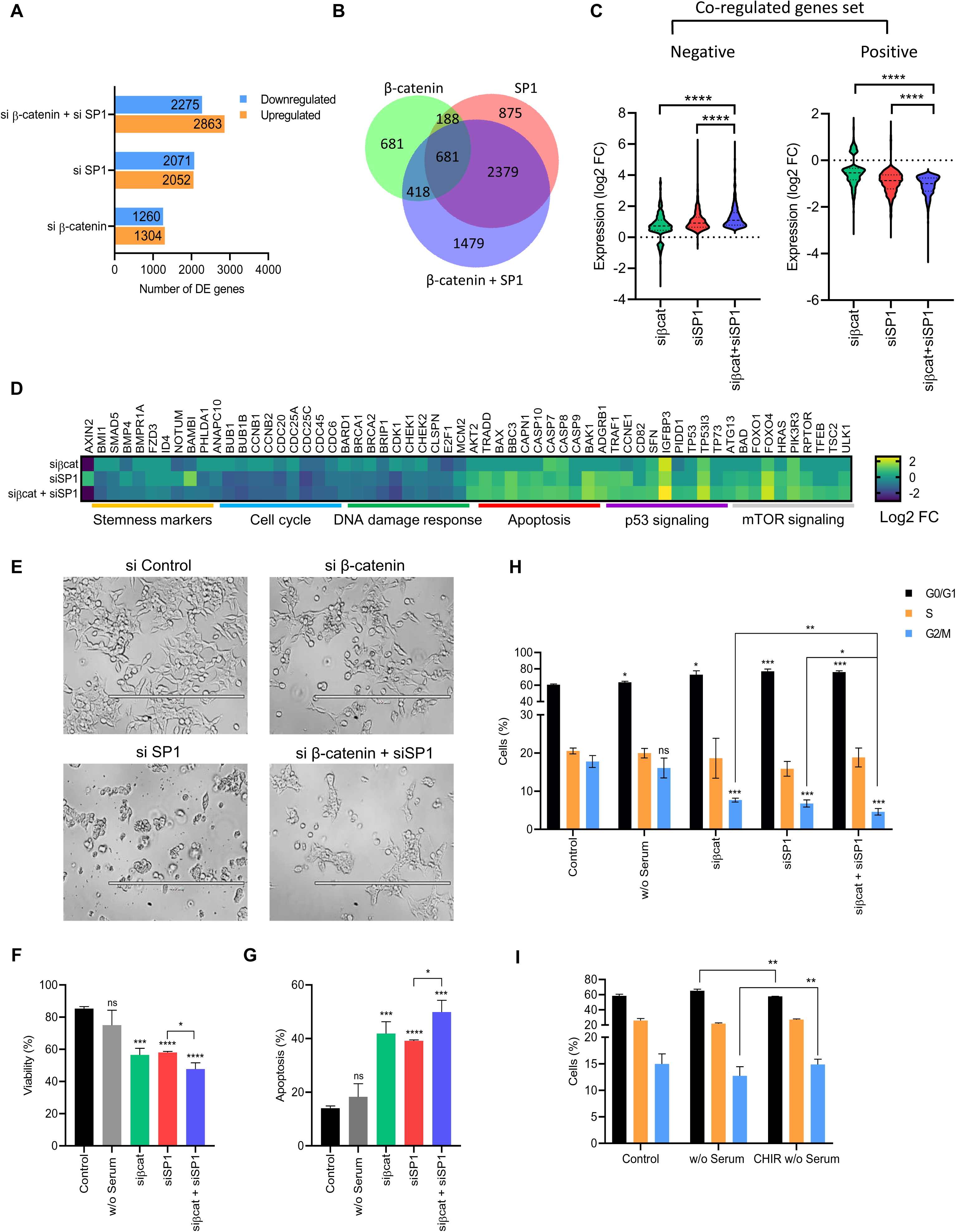
β-catenin and SP1 have synergistic transcriptional activity. **A.** Total number of differentially expressed genes in control siRNA, siβ-catenin, siSP1 and combination of both. **B.** Venn diagram showing overlap between the differentially regulated genes in cells treated with siβ-catenin only, siSP1 only and siβ-catenin + siSP1. **C.** Grouped log2 Fold change for common downregulated and upregulated genes upon SKD or DKD (Paired t-test performed to calculate significance). **D.** KEGG pathway enrichment analysis for the differentially expressed genes common in all three conditions. **E.** DIC images for cells treated with control siRNA, siβ-catenin, siSP1 and a combination of both. **F.** Quantification of viable cells after Annexin V/PI staining after treatment with siRNA against β-catenin and/or SP1. **G.** Quantification of apoptotic cells after Annexin V/PI staining after treatment with siRNA against β-catenin and/or SP1. Serum deprivation has been used as control. **H.** Cell-cycle analysis of cells treated with siRNA against β-catenin and/or SP1. Serum deprivation has been used as control. **I.** Cell-cycle analysis of cells treated with CHIR with serum deprivation. Serum deprivation has been used as control and unpaired t-test was performed to calculate significance for all.

Gene ontology (GO) analysis yielded downregulated genes enriched for cell proliferation and DNA replication, whereas upregulated genes were enriched for cell death-related pathways (autophagy, P53, and TAP63 pathways) (Supplementary Figure S2M&N). KEGG pathway analysis suggested notable decrease in the activity of pathways associated with the cell cycle and DNA damage response. Furthermore, several marker genes for colorectal cancer stem cells (CCSC) displayed reduced expression (marker list obtained from Wang et al 2021^14^) (Figure 2D), whereas genes linked to apoptosis and the p53 pathway exhibited increased expression. Notably, we also detected an upregulation of mTOR signaling (Figure 2D). As mTOR is known to promote tumor growth and crosstalk with the Wnt pathway¹⁵, it is plausible that concurrent depletion of β-catenin and SP1 shifts the cellular balance toward heightened mTOR activity to compensate and sustain cell proliferation¹⁵.

To test if β-catenin and SP1 regulate cell proliferation while simultaneously inhibiting cell death pathways, we profiled cell viability in all three KD conditions with serum deprivation as a control. While serum deprivation does not necessarily induce cell death, it does arrest cellular proliferation^16^. As anticipated, there was no alteration in overall cell viability during serum deprivation. We observed reduced cell viability and increased apoptosis upon depletion of β-catenin and SP1, with the highest effect being observed in the case of the combinatorial knockdown (Figure 2F&G and Supplementary Figure 3A). Cell cycle analysis showed an augmented accumulation of the G0/G1 population and a reduced G2/M population upon KD (Figure 2H and Supplementary Figure 3D). Further, a significant reduction in protein levels was observed in DNA replication (PCNA) and repair (RAD51) factors (Supplementary Figure S3B). We also observed cleavage of PARP (a hallmark of apoptosis^17^) in all three KD conditions (Supplementary Figure S3B, right panel).

Additionally, the changes associated with KD conditions were also replicated in serum deprivation (Supplementary Figure S3B, left panel), including the decrease in β-catenin and SP1 proteins. One of the primary reasons for these changes could be the absence of MYC which is one of the immediate-early genes upon stimulation via serum mitogens. Ectopic expression of MYC alone is sufficient to initiate the cell cycle, even in the absence of serum or growth factors^18^. While MYC is one of the classical targets of the Wnt/β-catenin pathway, it is also known to act alongside SP1 for the expression of cell cycle-related genes^19,20^. We monitored the expression of MYC under all the KD conditions, and observed that the expression of MYC decreased in the case of DKD and β-catenin KD but not SP1 KD (Supplementary Figure S3C). a similar cell cycle experiment using serum deprivation along with CHIR treatment which inhibits the activity of GSK3β, thereby acting as a Wnt agonist^21^. As anticipated, the reduction in the G2/M population due to the absence of growth signals was rescued when the cells were treated with CHIR (Figure 2I). CHIR mediated GSK3β inactivation leads to stabilisation of multiple proteins, therefore, we monitored if the ectopic expression of β-catenin and SP1 drives cell growth in the absence of serum. Although there was no discernible difference in the cell cycle profile but we did observe higher cell confluency in the cultures upon overexpression of β-catenin and SP1 (Supplementary Figure S4A&B). We also identified an overlap of 267 out of the 590 genes (∼45%) that are known to exhibit serum response (comprising both Myc-dependent and -independent serum response factors)^18^ (Supplementary Figure S4C). Collectively, these observations suggested that β-catenin and SP1 collaborate to enhance the expression of stemness markers, facilitating self-renewal in cells. They also support proliferative programs autonomously while hindering cell death, which can contribute to the maintenance of the CCSC population and overall increase in tumor growth.

### β-catenin and SP1 show chromatin co-occupancy at promoter and enhancer regions

To further investigate the mechanism via which β-catenin and SP1 act together at the level of chromatin., we examined the genome-wide chromatin occupancy profiles of β-catenin and SP1 by utilising the ChIP-seq datasets for β-catenin and SP1 in HCT116 cells^22,23^. As TCFs are the primary DNA-binding partners for β-catenin, we also included TCF7L2 into this analysis^23^. Among all TCF/LEF members, TCF7L2 emerged as highly expressed while TCF7L1 and TCF7 displayed lower levels of expression and LEF1 did not show expression (Supplementary Figure S5A). A total of 5383 peaks were called for β-catenin, 6362 peaks for TCF7L2 and 19,673 peaks for SP1. Since motifs for SP1 are much more abundant in the genome, therefore, the number of peaks is usually higher as compared to other transcription factors. Next, we overlapped these peaks to find the shared binding regions between the three proteins. Surprisingly, β-catenin shared ∼86% (4631) of its peaks with SP1 as opposed to ∼40% (2192) of its peaks with TCF7L2. Out of these, only a few were found to be devoid of SP1 occupancy (68 peaks) (Supplementary Figure S5B).

Further, we also incorporated the ChIP-seq profiles for H3K4me1 (mark for enhancers), H3K4me3 (mark for promoters), H3K27ac (mark for active chromatin regions) and RNA Polymerase II 5S (an indicator for active transcription) (ENCODE Project Consortium, 2012^23^) to characterise genomic regions. two distinct patterns of co-occupancy for β-catenin with SP1 and TCF7L2. In active enhancer regions (Cluster3: C3), we observed co-occupancy of β-catenin, TCF7L2, and SP1. While, on promoter regions (Cluster2: C2) only β-catenin and SP1 exhibited co-occupancy, while enrichment for TCF7L2 was limited (Figure 3A&B).

**Figure 3:**
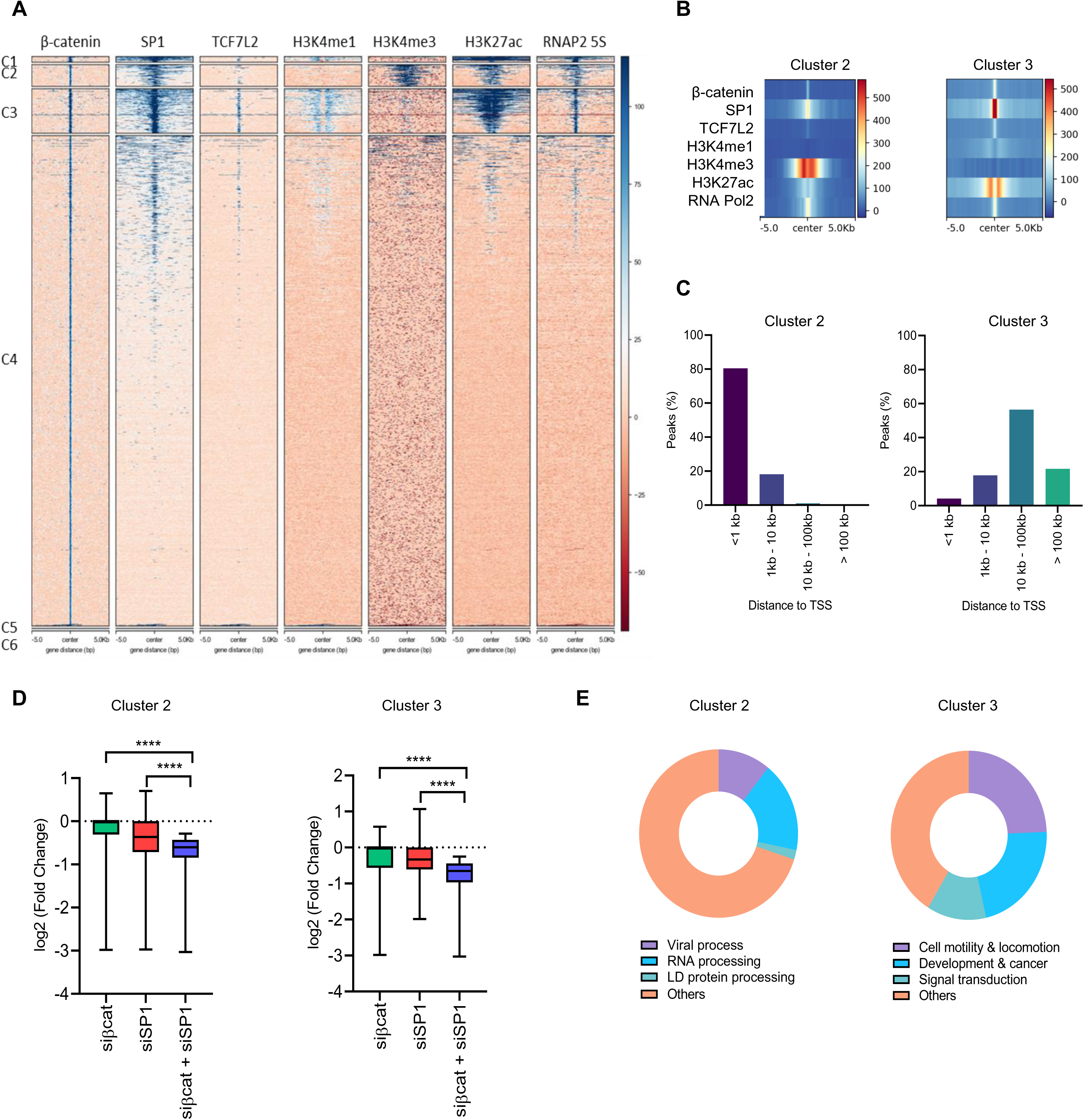
β-catenin and SP1 show chromatin co-occupancy at promoter and enhancer regions. **A.** Heatmaps for the ChIPseq profiles of β-catenin, SP1, TCF7L2, H3K4me1, H3K4me3, H3K27ac and RNA PolII 5S (Datasets: SRA012054, ENCSR000BMK, ENCSR000BSF, ENCSR000BVT, ENCSR161MXP, ENCSR333OPW, ENCSR661KMA, ENCSR000EUV). **B.** Peak intensities for all the ChIPseq datasets in Cluster 2 (C2: left) and Cluster 3 (C3: right) **C.** Percentage of peaks in C2 (left) and C3 (right) according to their distance to the nearest gene. **D.** Grouped log2 Fold change for common downregulated genes in C2 (left) and C3 (right) upon SKD or DKD (Paired t-test performed to calculate significance). **E.** GO analysis for common downregulated genes in C2 (left) and C3 (right) upon SKD or DKD.

Next, we plotted the distance of these peaks to the nearest transcription start site (TSS). For cluster 2 (C2), the majority of the peaks fell within less than 1 Kb to the nearest TSS, whereas for cluster (C3), the peaks exhibited a broader distribution, with most of them being situated at distances greater than 1 Kb to the nearest TSS (Figure 3C). These observations further confirmed the presence of promoters in C2 and enhancers in C3. To integrate the data from the RNAseq and the ChIPseq, We overlapped the genes obtained through these two methods and used the direct targets obtained for further analysis. We plotted the expression change for these direct targets upon the three KD conditions used earlier. Interestingly, both C2 and C3 genes showed the highest changes in the expression upon DKD as compared to any of the SKD conditions (Figure 3D). GO analysis of C2 and C3 associated genes showed the majority of C2 genes belonged to RNA and protein processing while C3 genes were involved in signal transduction,cancer and development-related processes (Figure 3E). These results indicate that SP1 is involved in driving the expression of β-catenin direct target genes, which may or may not be dependent on TCF7L2. These findings also indicated a dosage-dependent regulation, suggesting that SP1 and β-catenin alone may be adequate to stimulate the expression of their shared targets, but the presence of both leads to higher transcriptional activity. Furthermore, there appears to be a functional differentiation among the target genes they control. The genes dependent on TCF7L2 (C3) are primarily enriched for canonical Wnt targets, while the TCF7L2-independent genes (C2) are predominantly involved in RNA processing, which adds to the understanding of context-specific interactions involving β-catenin.

### SP1 co-localizes with the components of the Wnt-enhanceosome

The transcriptional response upon Wnt activation requires the assembly of the multi-subunit Wnt enhanceosome, comprising of β-catenin, TCF/LEF transcription factors, chromatin modifiers like CBP/p300, Pygo1/2, ChILs complex (LDB1, SSDP), BAF complex (SMARCA2/4, AIRD1A/1B/5B), among others^24^. Hence, we examined whether SP1 shares genomic binding sites with the components of the Wnt enhanceosome. To do so, we employed publicly available datasets for two of these components, SMARCA4 and LDB1^25,26^. We used peaks for β-catenin as the reference and aligned the ChIPseq datasets for SP1, TCF7L2, SMARCA4 and LDB1. We observed co-occupancy of all these proteins except LDB1 (Figure 4A&B). This indicates that SP1 might be a part of the Wnt enhanceosome itself. We checked the literature for known direct interactions between SP1 and Wnt enhanceosome components and found that SP1 can interact with EP300, LDB1 and SMARCA4. The interaction network made using SP1 and all the components of the Wnt enhanceosome showed how SP1 and β-catenin can be engaged with the rest of the factors^27^(Supplementary Figure 5C). To investigate whether the expression of any of the regulators for the transcriptional activity through the WREs is dependent on either β-catenin or SP1, we leveraged mRNAseq data to examine transcript levels for all the components of the Wnt enhanceosome. Our analysis revealed a significant reduction at the transcript levels for PYGO1, PYGO2, ARID5B, BCL9, TCF7L2 and HDAC1 (Supplementary Figure S6A-R). Notably, HDAC1, which typically functions as a repressor for WREs, was found to be under the positive regulation via β-catenin and SP1 indicating a feedback mechanism (Supplementary Figure S6R). While we observed occupancy of β-catenin and SP1 for nearly all the components we assessed, their perturbation did not lead to any significant change at the transcript levels (Supplementary Figure S7). Surprisingly, when we checked the protein expression for some of the components under β-catenin and SP1 KD conditions, reduced protein expression was observed for all the components analysed, including, BRG1, p300, CBP, and TCF7L2 (Figure 4E). Collectively, these findings suggest that SP1 engages with the constituents of the Wnt enhanceosome, as evidenced by their co-localization at shared target sites and their physical interactions. Moreover, both β-catenin and SP1 directly and indirectly regulate the expression levels of multiple Wnt enhanceosome components. Consequently, it can be inferred that the presence of these factors is crucial for the expression, and consequently, the assembly of the Wnt enhanceosome.

**Figure 4:**
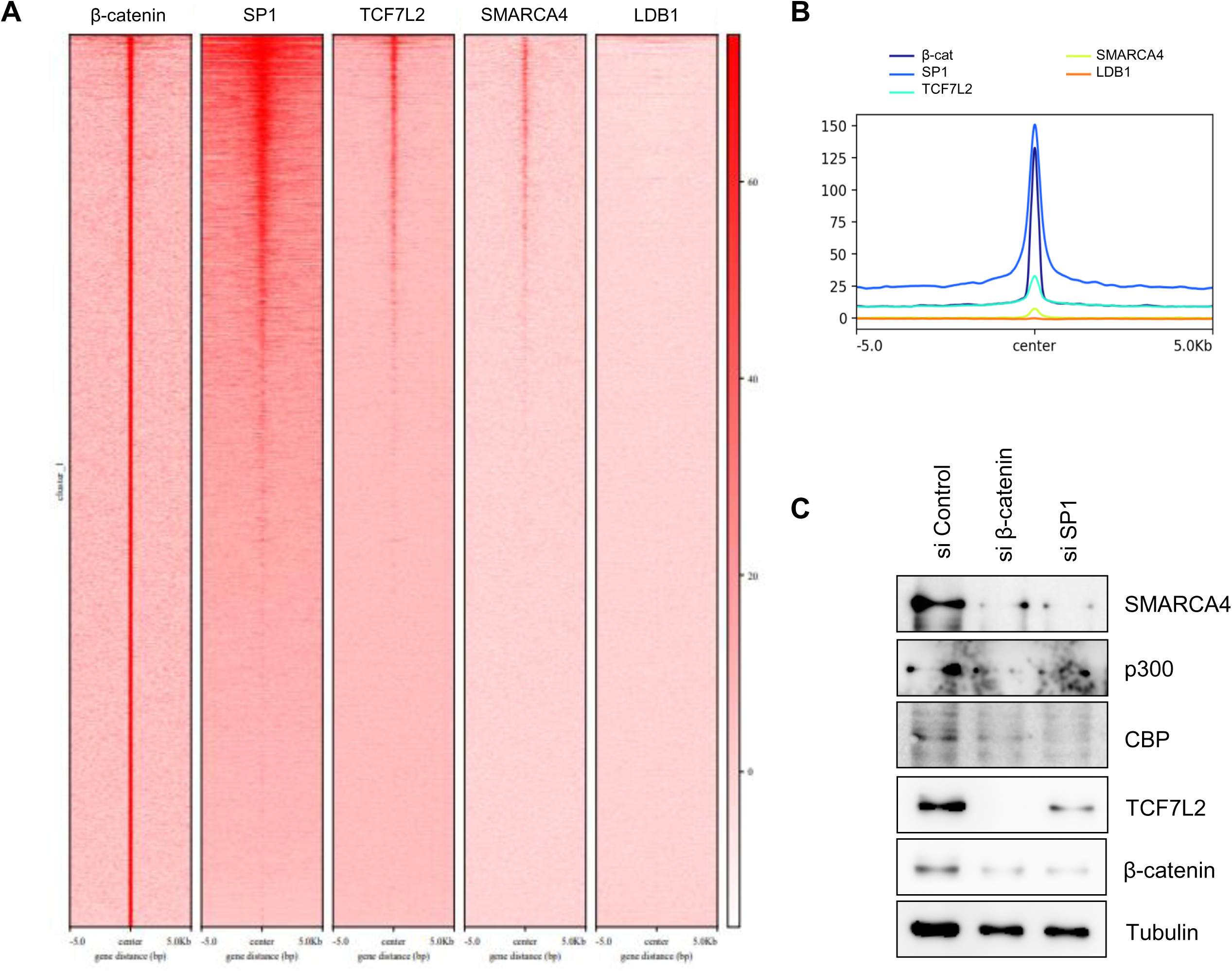
SP1 co-localises with the components of the Wnt-enhanceosome. **A.** Heatmap for ChIPseq for β-catenin, SP1, TCF7L2, BRG1 and LDB1. **B.** Peak summit for ChIPseq for β-catenin, SP1, TCF7L2, BRG1 and LDB1. **C.** WB image for BRG1, p300, CBP, TCF7L2 and β-catenin under siβ-catenin and siSP1 conditions.

### β-catenin and SP1 regulated genes contribute towards colorectal cancer development

β-catenin and SP1 are well-established to be upregulated in multiple cancers, and their high expression is associated with reduced patient survival. We overlapped the differentially expressed (DE) genes from our mRNAseq data with the DE genes derived from patient samples of colorectal adenocarcinoma (COAD) (Tang, Cho and Wang, 2022). We observed a substantial overlap between the two sets (Representation factor: 1.2; p < 2.960e-12) (Figure 5A). Further, the genes under the negative control of β-catenin and SP1 displayed no significant change between the normal and tumor samples (Figure 5B, left panel). However, genes under positive regulation displayed significant upregulation in the case of COAD (Figure 5B, right panel). These genes exhibited a strong enrichment for biological functions associated with the cell cycle and DNA replication, consistent with our previous observations (Figure 5C). Additionally, we assessed the impact of these shared targets on overall patient survival, revealing that several factors were negatively correlated with survival (Figure 5D). In summary, we established that the genes targeted by β-catenin-SP1 contribute to the development of CRC by promoting cell growth, consequently exerting a detrimental effect on patient survival.

**Figure 5:**
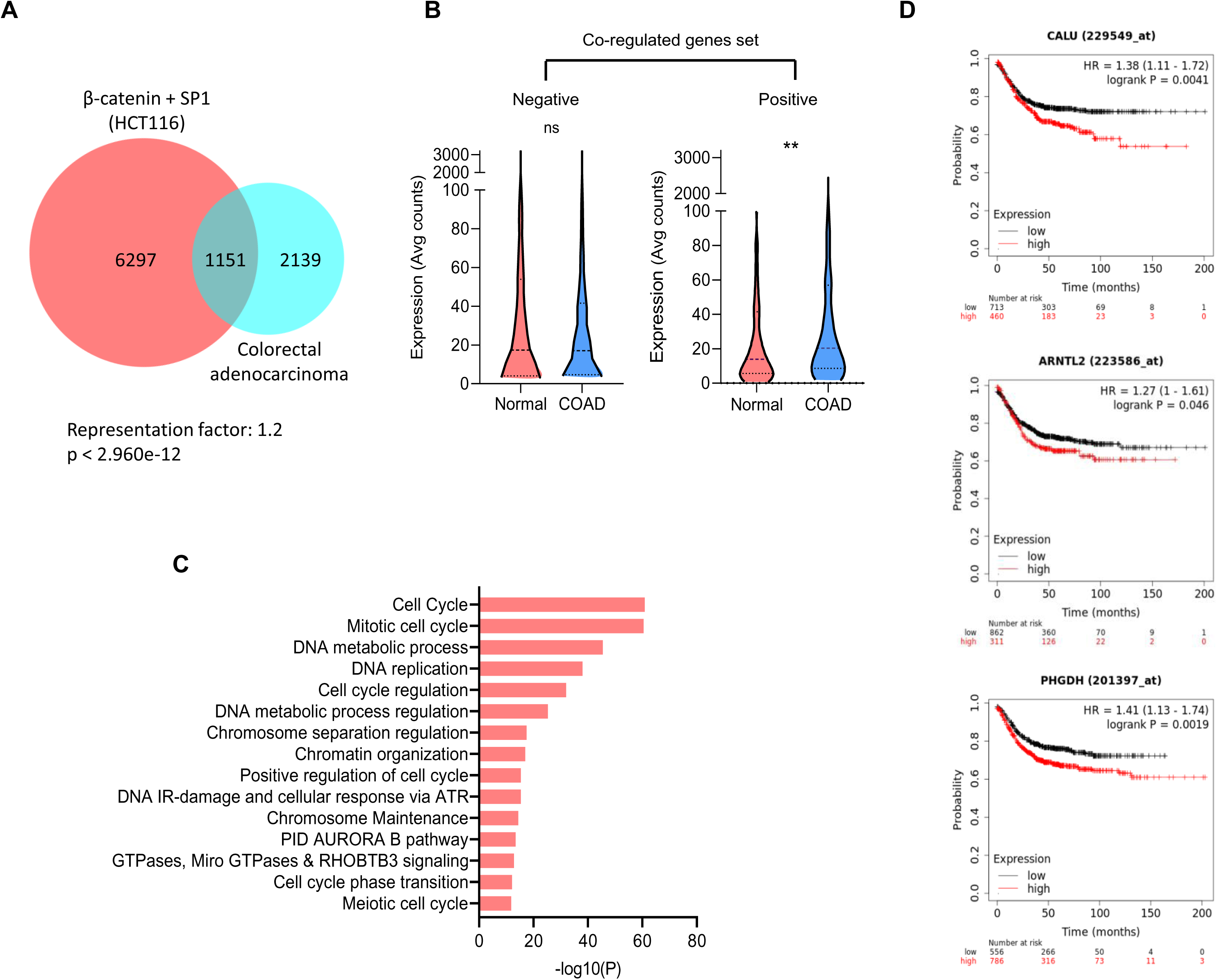
β-catenin and SP1-regulated genes contribute towards colorectal cancer development. **A.** Overlap between β-catenin and SP1 co-regulated genes obtained from mRNAseq in HCT116 and DE gene set for Colorectal Adenocarcinoma patient data from OncoDB using Tang et al., 2022. **B.** Expression counts for genes regulated by β-catenin and SP1 and overlapped with the COAD dataset (Paired t-test was performed to calculate significance). **C.** GO analysis for genes positively regulated by β-catenin and SP1 and overlapped with the COAD dataset. **D.** KM plots for selected targets (CALU: Calumenin; ARNTL2: Aryl Hydrocarbon Receptor Nuclear Translocator Like 2; PHGDH: Phosphoglycerate Dehydrogenase).

### Combinatorial chemical inhibition of β-catenin and SP1 promotes cell death and inhibits tumorigenesis

To assess the impact of β-catenin and SP1 on tumorigenesis within an in vivo context, we performed a cell-derived xenograft assay in mice. Following the induction of tumors, the mice were subjected to treatment with established inhibitors specific to either β-catenin, SP1, or both factors concurrently (Figure 6A). PNU 74654 (PNU) was employed as an inhibitor to impede the activity of Wnt/β-catenin signaling by obstructing the interaction of β-catenin and the TCFs^28^. Mithramycin-A (MT-A) was used to hinder the transcriptional activity of SP1 by binding to the GC-rich regions of the DNA^29,30^. Our findings revealed that mice treated with both inhibitors concurrently exhibited reduction in tumor growth compared to those treated with either inhibitor alone (Figure 6B). This reduction was evident in the overall tumor volume and weight suggesting synergistic effects of β-catenin and SP1 in the context of tumor development. (Figure 6 C&D). It is important to note that the frequency and the dosage of the inhibitors used were lower than those reported in the literature^31,32^. The reason for reducing the dosage was to mitigate potential toxicity to the animals.

**Figure 6:**
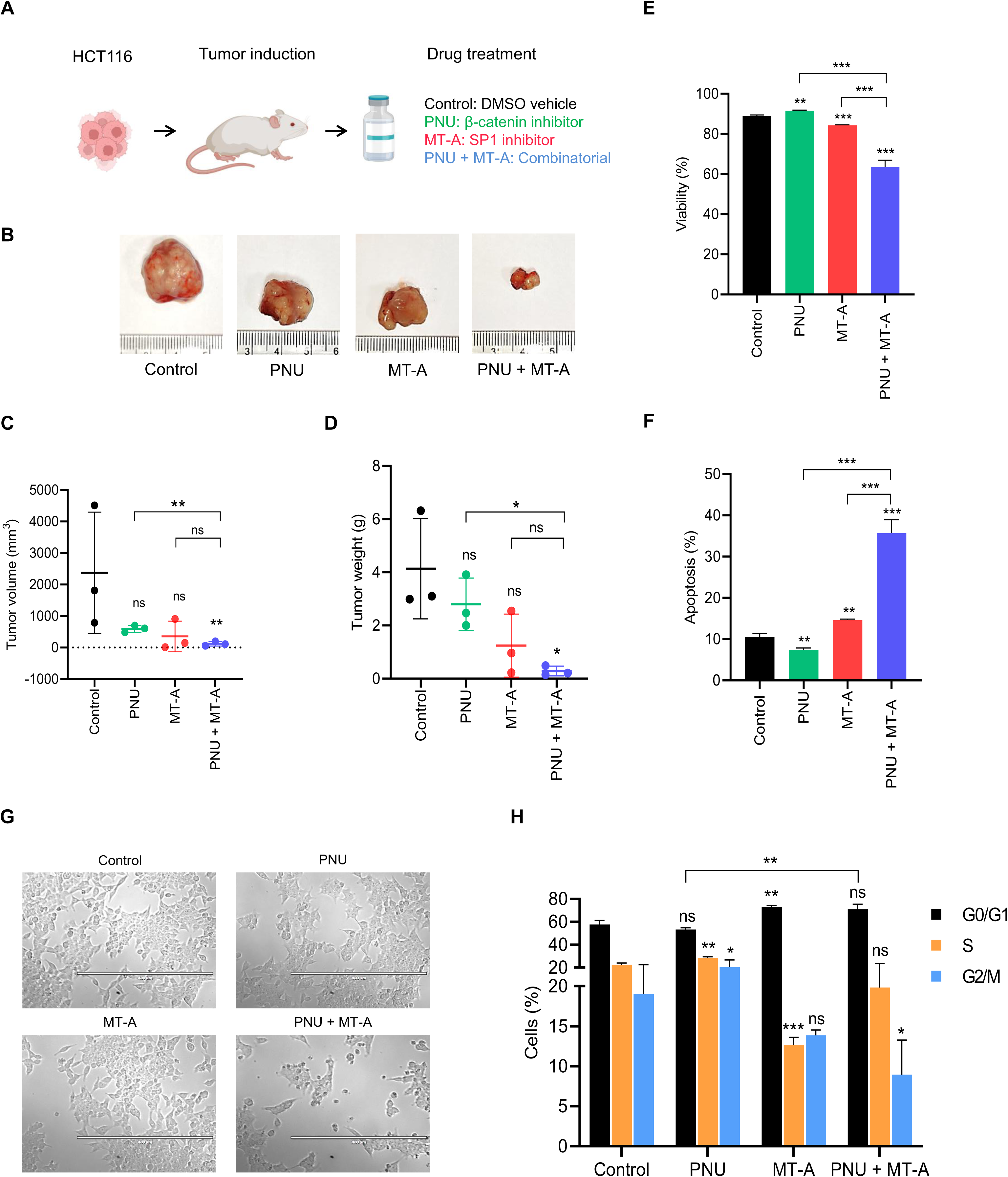
β-catenin and SP1 synergize to promote tumorigenesis. **A.** Schematic representation of the experimental design followed for the cell-derived xenograft study in mice. **B.** Representative images for the tumors harvested at the end of the treatments. **C.** Tumor volumes measured at the end of the treatments. Mean values: Control: 2371 mm3; PNU: 593.6 mm3; MT-A: 355.5 mm3; PNU + MT-A: 120.1 mm3. (Unpaired t-test used to calculate significance). **D.** Tumor weights measured at the end of the treatments. Mean values: Control: 4.137 g; PNU: 2.794 g; MT-A: 1.244 g; PNU + MT-A: 0.2844 g. (Unpaired t-test was used to calculate significance). **E.** Quantification of viable cells after Annexin V/PI staining after treatment with PNU, MT-A and PNU + MT-A (Unpaired t-test was used to calculate significance). **F.** Quantification of apoptotic cells after Annexin V/PI staining after treatment with PNU, MT-A and PNU + MT-A (Unpaired t-test was used to calculate significance). **G.** DIC images for cells treated with PNU, MT-A and PNU + MT-A. **H.** Cell-cycle analysis of HCT116 cells treated with PNU, MT-A and PNU + MT-A (Unpaired t-test was used to calculate significance).

Building upon these findings, we extended our investigation using the same treatment regime to assess the impact on cell survival and the cell cycle. Upon comparing the cell viability and the extent of apoptosis across all conditions, we observed a consistent pattern where the combinatorial treatment had a significantly more pronounced effect than single treatment (Figure 6E-G and Supplementary Figure S8A&B). Additionally, we examined the impact on the cell cycle profile and once again noted a drastic reduction in the G2/M population following the combinatorial treatment (Figure 6H and Supplementary Figure S8C).

### β-catenin and Sp1 overexpression leads to developmental defects in zebrafish

We aimed to investigate whether the β-catenin:Sp1 complex plays a role in development, using zebrafish development as our model system. However, information regarding Sp1 in zebrafish is relatively limited. The first study indicated that zebrafish Sp1 possesses conserved Zn-finger motifs and can induce transcription when expressed in a mammalian system^33^. Another study by our lab characterizes the expression and fundamental role of Sp1 during zebrafish development. Notably, zebrafish Sp1 exhibits localized expression in the anterior region and the gut in the larval stages. The target genes of Sp1 primarily regulate the cell cycle and cell death (unpublished work). Collectively, this demonstrated that zebrafish Sp1 fulfills the same fundamental functions as the human/mouse SP1 and is, therefore, a suitable model for studying its role during development. We applied the same treatment approach as previously illustrated (Figure 7A). Briefly, we titrated doses of both ctnnb1 and sp1 mRNA to identify doses that did not result in significant deformities or mortality. Subsequently, we injected *ctnnb1* and/or *sp1* mRNA in zebrafish at 1 cell stage and monitored the degree of mortality and phenotypic deformities until 2dpf. During the early stages, there were slight developmental delays associated with the overexpression of *ctnnb1* and *sp1*, but most of these delays resolved after the Shield stage (6 hpf, i.e., post-gastrulation). It is known that overexpression of *ctnnb1* leads to anteriorisation due to its role in anteroposterior axis formation^34^. Interestingly, we observed similar defects in the case of *sp1* overexpression, suggesting their co-involvement in some developmental processes, if not all (Figure 7B&C). Notably, there was a significant increase in the number of larvae displaying defects in their posterior structure (Figure 7C). Quantification of mortality and deformities further revealed that the most pronounced effects were observed in the case of double overexpression (Figure 7D&E). The proliferation marker mki67 exhibited the highest expression levels when both β-catenin and Sp1 were overexpressed at 10hpf (Figure 7F). These observations strongly suggest that β-catenin and Sp1 jointly participate in specific developmental processes, such as cell proliferation and anterior-posterior axis formation.

**Figure 7:**
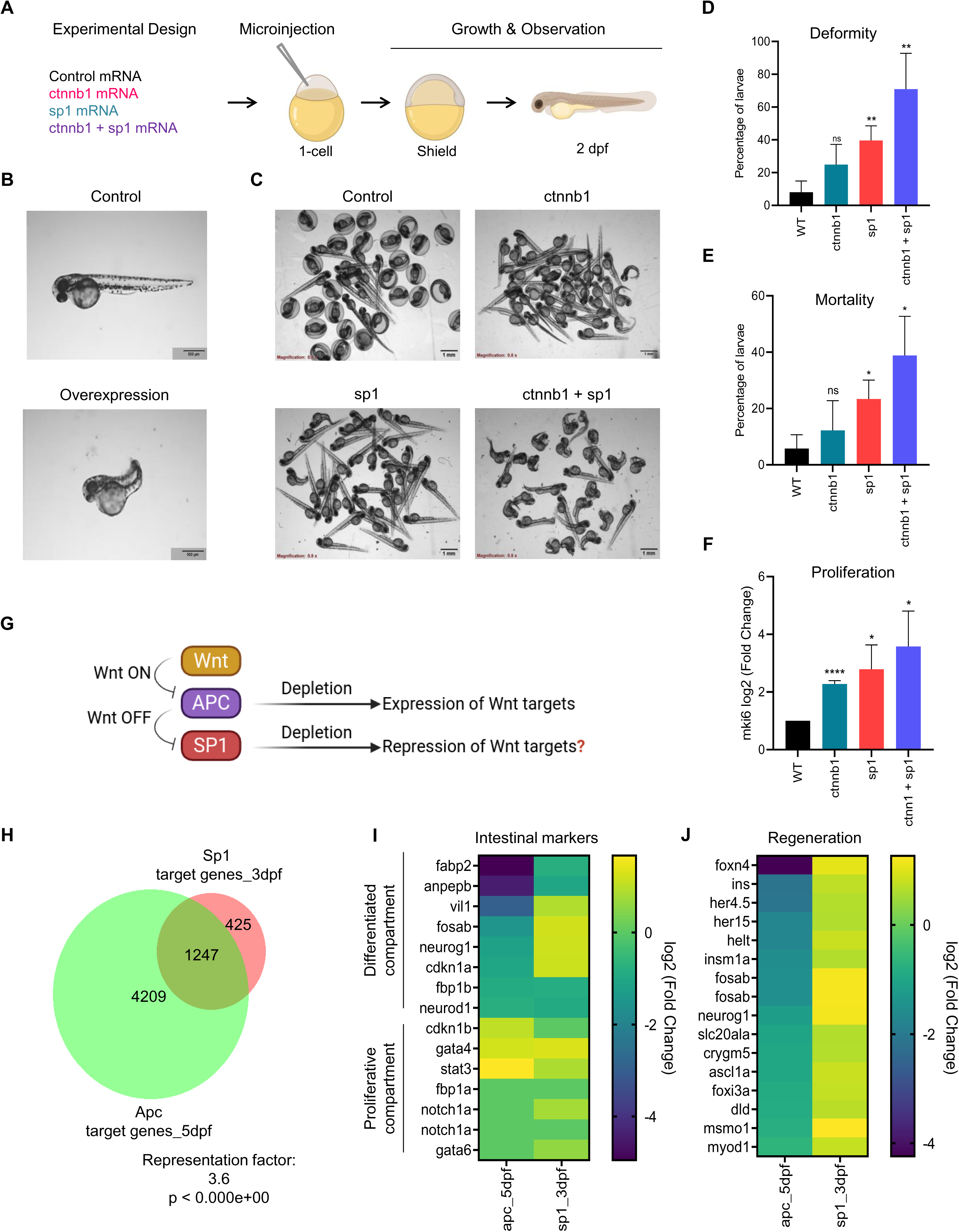
Overexpression of β-catenin and SP1 leads to similar developmental defects in zebrafish. **A.** Schematic representation of the experimental design followed for the overexpression study in zebrafish. **B.** Representative higher magnification images for zebrafish for control and overexpression phenotype at 2dpf. Magnification = 2X, Scale bar = 500 µm. **C.** Representative brightfield images for zebrafish injected with control (GFP), ctnnb1 mRNA, sp1 mRNA or both at 2dpf. Magnification = 0.8X, Scale bar = 500 µm. **D.** Quantification of larvae with deformities at 2dpf (Unpaired t-test was used to calculate significance). **E.** Quantification of mortality at the end of 2dpf (Unpaired t-test was used to calculate significance). **F.** Expression values of mki67 using qRT-PCR at Bud (10hpf) stage (Unpaired t-test was used to calculate significance). **G.** Schematic representation of the hypothesis for the contrasting actions of Sp1 and Apc in zebrafish. **H.** Venn diagram depicting the overlap between the DE genes obtained after treatment with *sp1* morpholino until 3dpf or *apc* deletion at 5dpf. **I.** Expression values for the intestinal epithelial markers (differentiated and stem cell markers) in *apc*^-/-^ line at 5dpf and *sp1* morpholino treatment until 3dpf. **J.** Expression values for the intestinal regeneration markers in apc-/- line at 5dpf and *sp1* morpholino treatment until 3dpf.

Next, we integrated another major component of the Wnt signaling pathway in our study-Apc. Apc is a negative regulator of the Wnt pathway and promotes the degradation of β-catenin^35^. We obtained the RNA isolated from *apc* mutants and WT siblings at 5 dpf. We used these RNA samples to perform mRNAseq. Since Apc is a negative regulator of the Wnt pathway, depletion of Apc should result in the activation of Wnt target genes. Based on our hypothesis, if Sp1 collaborates with Wnt signaling during zebrafish development, then Sp1 depletion should result in the repression of Wnt targets (Figure 7G). To check whether the reduced levels of Apc or Sp1 affect the same set of genes (potential Wnt responsive genes), we overlapped the DE genes obtained from both datasets. Strikingly, ∼74% of Sp1 targets were found to be affected by *apc* deletion as well (Representation factor: 3.6; p < 0.0001) (Figure 7H). We proceeded with this common set of genes and analyzed the expression of a few relevant factors in both the depletion conditions. We noticed that the profiles obtained under both these conditions were completely contrasting to each other, aligning with our hypothesis. Several factors that are enriched in the differentiated compartment of the zebrafish gut epithelium exhibited an increase in the case of Sp1 depletion and a decrease with *apc* deletion. In contrast, certain markers associated with the proliferative compartment showed downregulation in response to Sp1 depletion, with relatively little change in *apc* mutants (Figure 7i). However, we observed a consistent pattern in the case of gut regeneration markers. The expression of all regeneration markers was elevated in *apc* mutants but reduced in Sp1 morpholino-injected larvae (Figure 7J). In summary, our findings reveal the opposing roles of Apc and Sp1 during zebrafish development, hinting towards a potential collaboration between Sp1 and Wnt signaling, facilitating the continuous regeneration and growth of the gut epithelium.

## Discussion

β-catenin functions as a central transcriptional co-activator in the canonical Wnt signaling pathway and engages with a diverse set of transcriptional regulators to fine-tune gene expression in a context-dependent manner. This regulatory flexibility enables Wnt signaling to execute diverse biological outcomes during development, differentiation, and the maintenance of tissue homeostasis^1^. However, the precise mechanisms and functional implications of β-catenin’s transcriptional partnerships remain incompletely understood. In this study, we identify SP1 as a critical interactor of β-catenin that promotes its stability during Wnt activation and facilitates target gene transcription. Our findings uncover a cooperative role for SP1 and β-catenin in the regulation of cell cycle progression and apoptosis, and show that both factors can independently support cell growth, an attribute relevant to both developmental and pathological contexts.

The early expression of both SP1 and β-catenin, combined with the embryonic lethality observed upon their depletion^36–38^, points toward their essential role in driving rapid and autonomous proliferation during early embryogenesis. We propose that the β–catenin–SP1 complex forms part of a core transcriptional program that governs this growth trajectory. Similarly, in cancer, particularly during the early phases of tumor initiation, uncontrolled proliferation precedes later hallmarks such as epithelial-to-mesenchymal transition and chemoresistance. The ability of SP1 and β-catenin to promote autonomous growth likely contributes to the hyperproliferative state characteristic of early tumorigenesis.

At the chromatin level, we speculate that SP1 and β-catenin co-occupy DNA regions both in the presence and absence of TCF7L2. Intriguingly, sites co-bound by β-catenin, SP1, and TCF7L2 are enriched for enhancer-associated chromatin marks, while β-catenin-SP1-only loci display promoter-associated features. These findings suggest a division of labor, where SP1 preferentially binds promoter regions and may recruit β-catenin, consistent with prior reports^39^, while β-catenin may similarly enhance SP1’s occupancy at distal Wnt-responsive enhancers. The combinatorial and context-specific recruitment of SP1 and β-catenin likely determines the transcriptional output of Wnt signaling. In cancer cells, where these factors are highly expressed, this regulatory balance may be perturbed, leading to pervasive and promiscuous binding across both promoter and enhancer elements. Time-resolved binding studies following Wnt stimulation will be crucial to understand the dynamics of SP1 and β-catenin recruitment and their contribution to gene expression kinetics.

Importantly, we also identify SP1 as a potential component of the Wnt enhanceosome. We show that SP1 co-occupies genomic loci with core enhanceosome factors, including BRG1 (SMARCA4), and regulates their expression at both transcript and protein levels. Although we have not explicitly tested whether SP1–BRG1 co-occupancy is restricted to TCF7L2-bound sites, our analysis indicate the existence of β–catenin–SP1 loci lacking BRG1, possibly reflecting alternative modes of transcriptional activation at promoter-proximal sites. Furthermore, SP1’s known ability to dimerize and facilitate DNA looping may contribute to long-range enhancer–promoter communication. This function parallels other SP family members such as SP5 and SP8, which are known to mediate chromatin looping in the context of Wnt-regulated transcription^9^. Together, these results suggest that SP1 contributes to Wnt-responsive gene regulation not only as a direct DNA-binding transcription factor but also as a structural and regulatory component of the Wnt enhanceosome.

Functionally, our study establishes that SP1 and β-catenin together promote cellular proliferation and suppress apoptosis. While this cooperation is important for organ growth and adult stem cell maintenance under physiological conditions, its overactivation leads to pathological outcomes, including tumor formation. The gene expression program co-regulated by SP1 and β-catenin is conserved across species and is highly enriched in CRC. Pharmacological co-inhibition of SP1 and β-catenin resulted in a synergistic reduction in tumor cell growth, indicating that therapeutic strategies targeting both factors simultaneously may be more effective than inhibiting either protein alone.

Beyond cancer, the parallels between developmental and tumorigenic gene expression programs are increasingly recognized. Reactivation of developmental pathways such as Wnt, Notch, and Hedgehog is a well-established feature of cancer progression, contributing to processes such as epithelial–mesenchymal transition (EMT), angiogenesis, and cancer stem cell maintenance^40^. Within the intestinal epithelium, Wnt signaling is a key driver of endoderm specification during development and intestinal stem cell (ISC) maintenance during adult tissue homeostasis. Dysregulation of this pathway is a hallmark of CRC initiation and progression^41^. In light of this, we explored whether the SP1–β–catenin gene signature observed in cancer is also relevant during embryonic development, using zebrafish as a model system. Previous work from our lab has shown that zebrafish Sp1 is ubiquitously expressed during early embryogenesis. Eventually, it becomes increasingly restricted to the head and gut regions, indicating a potential role in neural and endodermal development. Additionally, Sp1 regulates both cell proliferation and endoderm formation in zebrafish embryos, mirroring its role in human cells. When overexpressed individually or in combination, Sp1 and β-catenin induced anteriorization phenotypes in zebrafish embryos. This is consistent with the known function of Wnt/β-catenin signaling in anterior–posterior axis formation, where excess β-catenin leads to anterior structure expansion and posterior truncation^42^. The fact that Sp1 overexpression elicited similar defects suggests that it may also participate in axis specification. The increased penetrance and severity of phenotypes observed upon co-overexpression of Sp1 and β-catenin further support the notion that these factors function synergistically during early development.

Finally, our work uncovers an antagonistic regulatory relationship between Sp1 and Apc, a well-characterized negative regulator of Wnt signaling. A significant fraction of their target genes overlap, particularly those involved in intestinal regeneration, but are regulated in an opposite manner. This antagonism points to a fine balance between positive and negative inputs required to maintain gut epithelial turnover, with potential implications for both development and disease.

In summary, we delineate a conserved β–catenin–SP1 transcriptional module that plays a critical role in early development, tissue homeostasis, and colorectal tumorigenesis. Our findings position SP1 as both a transcriptional and structural component of the Wnt enhanceosome, capable of modulating enhancer–promoter communication and gene expression output in a context-dependent manner. The functional cooperativity between SP1 and β-catenin underscores the importance of transcription factor partnerships in the regulation of complex biological programs and provides a strong rationale for targeting this axis in Wnt-driven cancers (Figure 8).

**Figure 8.**
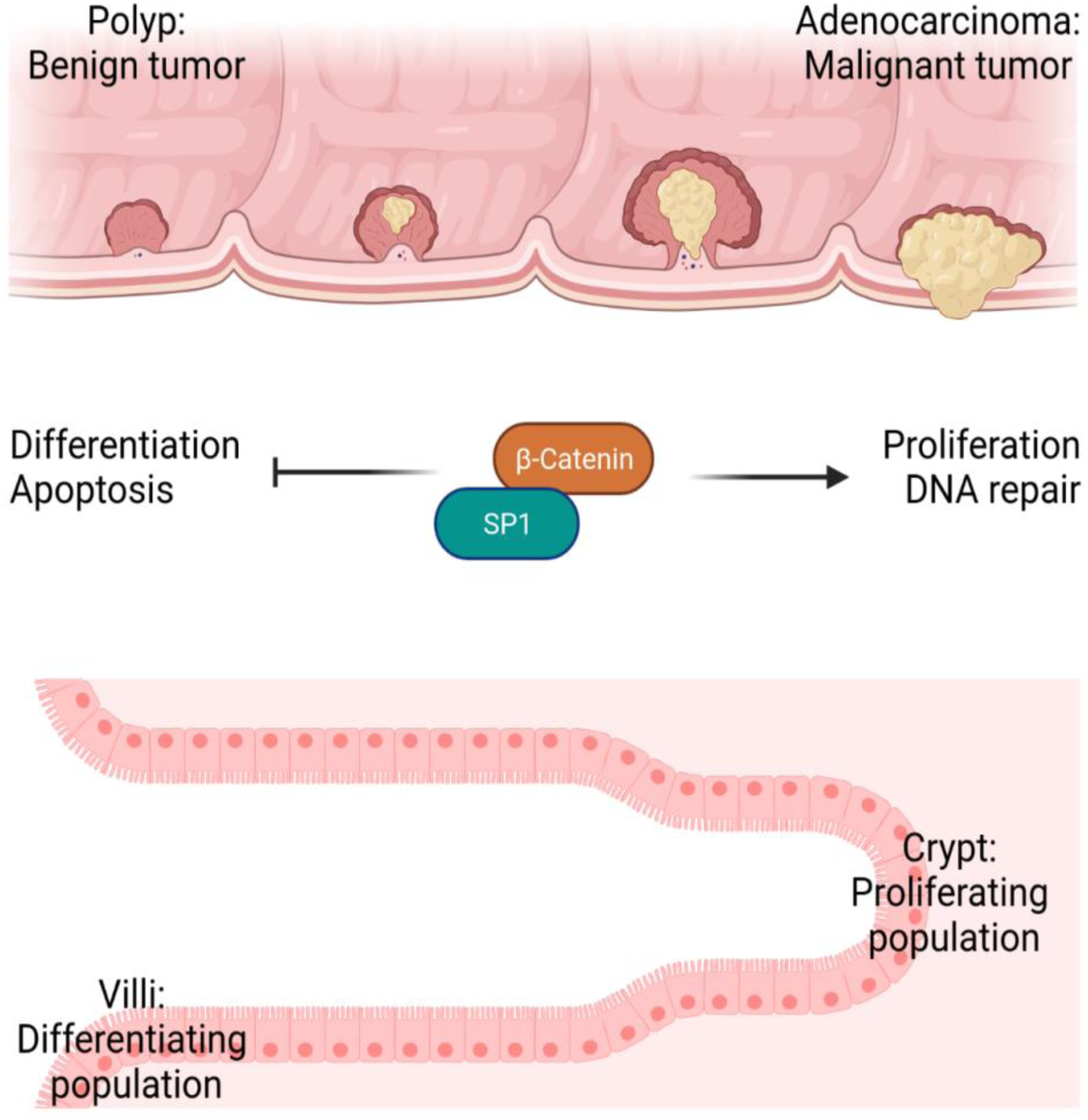
Model for the conserved function of the β-catenin and SP1 complex in development and disease. β-catenin and SP1 promote cellular proliferation and prevent cell death. During regular growth and development, it can help in organ growth and maintenance of adult stem cells (lower panel). However, in case of abnormal stimulation, it can contribute towards the aggressiveness of cancer.

### Limitations

We aimed to investigate whether the assembly of the Wnt enhanceosome is impacted in the absence of SP1, while simultaneously restoring β-catenin levels through ectopic expression. However, given that the depletion of SP1 resulted in the reduction of multiple components of the Wnt enhanceosome, we were unable to address this aspect. Examining the role of SP1 in the assembly of the Wnt enhanceosome in various contexts is of utmost importance. Additionally, future studies should delve into the regulatory role of SP1 concerning Wnt target genes in the absence of TCFs. It would be intriguing to explore the impact on the expression of C2 and C3 genes when TCF7L2 is removed from the system.

## Materials and methods

### Human cell culture, transfection and chemical inhibitor treatment

HEK293T and HCT116 cells were obtained from the European Collection of Cell Cultures (ECACC). Cells were cultured in DMEM (Thermo Fisher Scientific) supplemented with 10% heat-inactivated Fetal Bovine Serum (Thermo Fisher Scientific) and 1% Anti-Bacterial, Anti-Mycotic (Thermo Fisher Scientific). Serum deprivation was performed by incubating the cells in serum free culture medium 48 h prior to harvesting. For transfections, 0.3 x 10^6^ cells were seeded in a 6-well culture dish 16 h prior to transfection. Each well was transfected with either 2 µg of plasmid or 10 pmol of siRNA using Lipofectamine 3000 or Lipofectamine RNAimax (Thermo Fisher Scientific) respectively, according to the protocol provided by the manufacturer. The cells were harvested 48 h post-transfection for RNA and protein isolation. A list of all the siRNAs and plasmids sequences used is provided in supplementary table 1 and 2, respectively. For treatment with chemical inhibitors in HCT116 cell culture, Mithramycin A was diluted to a concentration of 25 nM, PNU 74654 to 20 μM and CHIR to 3 μM in prewarmed complete DMEM before mixing and adding to the culture dish. The cells were replenished with fresh medium and inhibitor every 24 h until harvesting. The cells were treated for 48 h before harvesting. Details for the chemicals used is provided in the supplementary table 2.

### RNA isolation, mRNA sequencing and analysis

The cells/zebrafish embryos were lysed in 1 ml RNAiso Plus (Takara) followed by standard extraction protocol. Briefly, the cells/zebrafish embryos were homogenised using Pellet Pestle Motor Kontes. 200 μL chloroform was added followed by centrifugation at 12000 rpm at 4℃, 15 min . The aqueous layer was collected in a fresh tube, followed by another chloroform wash. Precipitation was performed by adding 10% of 3 M sodium acetate solution and an equal volume of 100% isopropanol containing Glycoblue coprecipitant (Invitrogen) and incubating at -20℃, overnight. RNA was pelleted at 12000 x g at 4℃ for 1 h, followed by two washes of 75% ethanol at 12000 x g, room temperature (RT), for 5 min. Total RNA was resuspended in nuclease-free water (Ambion) followed by quantification using Qubit RNA HS system (Thermo Fisher Scientific) and RNA integrity determination using RNA 6000 Nano Kit on Bioanalyzer 2100 (Agilent). RNA samples with RIN values greater than 8 were used for library preparation.

500 ng of total RNA was subjected to mRNA purification using NEBNext Poly(A) mRNA Magnetic Isolation Module (New England Biolabs) according to the manufacturer’s instructions. The purified mRNA was used for library preparation using the NEBNext Ultra II RNA Library Prep Kit for Illumina (New England Biolabs) using the protocol provided in the kit. The final libraries were purified using the HighPrep PCR Clean-up System (MagBio Genomics, USA). The libraries were quantified using the Qubit 1X HS DNA system (Thermo Fisher Scientific). All the libraries were pooled in equimolar ratios and subjected to 75 bp PE chemistry on Nextseq550 sequencer (Illumina) aiming at least 30 million reads per sample. The bcl files obtained from sequencing were demultiplexed and converted to fastq files for further analysis. The sequencing reads were trimmed using Trimmomatic-0.39 and aligned to GRCh38 genome assembly for humans using the HISAT2 alignment program^43,44^. Gene feature counts were calculated using the FeatureCounts package from Rsubread^45^. EdgeR was used to perform differential expression analysis for 3 replicates per condition^46^. Graphpad prism was used for creating the representative plots for the performed analysis. Metascape was used for performing the Gene Ontology analysis^47^.

### Protein extraction, western blotting and quantification

Protein extraction was performed by lysing the cells in RIPA buffer (50 mM Tris-HCl, pH 7.5, 150 mM NaCl, 1% Nonidet P-40, 0.5% sodium deoxycholate and 0.1% sodium dodecyl sulphate) for 30 min on ice followed by centrifugation at 20000 x g for 30 min, 4℃. The supernatant obtained was quantified using the Pierce BCA Protein Assay kit (Thermo Fisher Scientific). Total cell lysate (25 µg) was boiled in SDS loading dye for 5 min at 98℃ and loaded for western blotting unless mentioned otherwise. ImageJ software was used for densitometric analyses: https://lukemiller.org/index.php/2010/11/analyzing-gels-and-western-blots-with-image-j/. All values were normalised with the loading control, followed by internal normalisation by the control condition.

### Immunofluorescence imaging and quantification

For immunofluorescence assay, cells were cultured on fibronectin-coated glass coverslips and fixed using 4% paraformaldehyde (PFA) (Merck) in 1X PBS at room temperature (RT) for 15 min. Permeabilization was performed using 0.25% Triton X-100 for 5 min, followed by blocking in 10% bovine serum albumin (BSA) in 1X PBS at RT for 1 h. The cells were then incubated in primary antibody (1:100 in 5% BSA in 1X PBS) at 4°C for 16 h, followed by four washes in 1X PBS at RT, incubation in secondary antibody (1:1000 2% BSA) for 1 h, and four washes in 1X PBS. Imaging was performed using the Zeiss 710 LSM confocal microscope. Quantification of the nuclear intensity was performed using the ImageJ software. The nuclear area within each cell was selected using 4′,6-diamidino-2-phenylindole (DAPI) staining as the region of interest (ROI). Quantified nuclear intensity was normalized with the nuclear area and DAPI. At least 300 cells were used for quantification in each set. A list of the antibodies used for western blotting is provided in supplementary table 2.

### Analysis of chromatin immunoprecipitation (ChIP)-seq datasets

The following datasets were used for analysis: SRA012054, ENCSR000BMK, ENCSR000BSF, ENCSR000BVT, ENCSR161MXP, ENCSR333OPW, ENCSR661KMA, ENCSR000EUV (ENCODE Project Consortium, 2012). The sequencing reads were trimmed using Trimmomatic-0.39 and aligned to GRCh38 genome assembly for humans using the BWA alignment program^43,48^. Peak calling was performed using MACS2 with default parameters and q value 0.05^49^. Consensus peaks from the biological replicates were extracted using a custom R script from Roman Cheplyaka (https://ro-che.info/articles/2018-07-11-chip-seq-consensus). deepTools 3.3.2 and Integrative Genomics Viewer (IGV) were used to create the representative figures for the performed analysis^50,51^. BigWig files were generated using bamCoverage (deepTools). Peaks were annotated to the nearest gene using Homer and classified into promoter (±5 Kb) and non-promoter regions. Clusters were annotated using Homer. Gene ontology analysis was performed using Metascape and WebGestalt^47,52^.

### Annexin V/Propidium Iodide staining and quantification for cell viability and cell cycle analysis

Cell viability assay was performed using Annexin V/FITC kit (Abcam) according to the protocol provided by the manufacturer. Briefly, cells were trypsinized and washed with 1X PBS. The cells were resuspended in 250 μL 1X Annexin V binding buffer, followed by the addition of 2.5 μL Annexin V-FITC and 2.5 μL propidium iodide. The suspension was incubated in the dark for 5 min at RT. The Annexin V-FITC signal was then analysed via BD FACSCelesta™ Cell Analyzer (Ex=488 nm, Em=530 nm) using a FITC signal detector and a phycoerythrin emission signal detector to detect PI staining. The results were analysed using FlowJo™ v10.8.1 Software (BD Life Sciences). The debris was removed by defining an area of the FS vs. SS, followed by gating the single cell population using an FSC height vs. FSC Area plot. The singlet data was further classified as live (Annexin V-PI-), early apoptotic (Annexin V+PI), late apoptotic (Annexin V+PI+), or ‘necrotic’ (Annexin V-PI+) using a quadrant gate. Cell cycle assay was performed using propidium iodide staining (Merck). Cells were trypsinized and washed with 1X PBS, followed by fixing in ice-cold 70% ethanol overnight. The cells were subsequently washed with 1X PBS twice and incubated with 0.4 mg/ml RNase A overnight. The samples were then incubated with 50 µg/mL Propidium iodide for 5 min at RT. Data for the PI fluorescence was collected to a total count of 10,000 nuclei via BD FACSCelesta™ Cell Analyzer using phycoerythrin emission signal detection. The cell cycle fractions for G0/G1, S, G2/M were analysed using the BD FACSDiva Software program (BD Biosciences).

### Patient transcriptome data analysis

Gene counts for all the differentially expressed genes in colon adenocarcinoma were obtained from OncoDB (http://www.oncodb.org) with the parameters: FDR Adjusted p-value < 0.001, log2 Fold change > 1). Overlap between mRNAseq datasets was obtained using Venny 2.1.0 (https://bioinfogp.cnb.csic.es/tools/venny), and the Venn diagrams were generated using BioVenn (https://www.biovenn.nl/index.php). GO analysis for the common target genes was performed using the online version of Metascape^47^. Patient survival plots were generated using the online tool Kaplan-Meier Plotter (https://kmplot.com/analysis).

### Mice rearing and xenograft assays

NOD/SCID strain of mice was used for the study and housed in a pathogen-free facility according to the standard husbandry protocols at the National Facility for Gene Function in Health and Disease, IISER, Pune. For the cell-derived xenograft experiment, each NOD/SCID mouse was injected with 1 X 10^6^ HCT116 cells subcutaneously in a 1:1 mixture with Matrigel (Corning). At the end of 2 weeks post-injection, mice were treated with either 1X PBS or PNU 74654 (PNU) (0.15 mg/kg) or Mithramycin-A (MT-A) (0.5 mg/kg) or a combination of PNU (0.15 mg/kg) and MT-A (0.5 mg/kg) once a week for a total of 6 weeks. At the end of the experiment, measurements were taken for the tumor weight and volume. Tumor volume was calculated using the formula of 0.5 × length × width².

### Zebrafish husbandry, knockdown and overexpression assays

Wild-type strains of zebrafish, AB & TU; and transgenic sox17-eGFP lines were maintained according to the standard zebrafish husbandry protocols under the National Facility for Gene Function in Health and Disease, IISER, Pune. Embryos and larvae were grown at 28℃ in 1X E3 medium (5 mM NaCl, 0.33 mM CaCl_2_, 0.33 mM MgSO_4_, 0.17 mM KCl, and 0.1% Methylene Blue) till harvesting. Bright field and fluorescence imaging were performed using a stereo microscope (Olympus SZX7)

Morpholino for *sp1* and control morpholino were obtained from Gene Tools. Morpholinos were resuspended in nuclease-free water according to the instructions provided by the manufacturer. Embryos were injected with morpholinos at 0.5 nM concentration at 1 cell stage and grown till the indicated stage and harvested depending on the downstream application. Capped mRNA for Sp1 and β-catenin were synthesized using Ambion mMESSAGE mMACHINE™ SP6 Transcription Kit (ThermoFisher Scientific). mRNA was injected at a total concentration of 3 ng/µl at 1 cell stage and grown till the indicated stage and harvested depending on the downstream application. For quantification of deformity and mortality, 100 embryos were injected per condition per replicate. For RNA isolation, 10 embryos were harvested per replicate and processed as described previously. A list of the morpholino sequences and plasmids used for fish experiments are provided in the supplementary tables 1 and 2, respectively. Library preparation and data analysis for the mRNAseq performed for sp1 morpholino-injected larvae (3 dpf) and *apc^-^*^/-^ larvae (5 dpf) with their respective WT control siblings was performed according to the protocol described previously.

### Statistical Analysis

All experiments were performed in biological triplicates unless specified otherwise. Statistical analyses between two groups were performed using the two-tailed unpaired Student’s t-test. A confidence level of 0.05 was considered statistically significant unless stated otherwise.

## Data Availability

Sequencing data will be deposited in the Gene Expression Omnibus (GEO) and made publicly available upon peer-reviewed publication.

## Declarations Funding

The work was supported by research grant from Department of Biotechnology (DBT), Government of India to KS and SG (BT/PR26289/GET/119/226/2017). SG is also a recipient of the JC Bose Fellowship (JCB/2019/000013) by the Science and Engineering Research Board, Government of India. We thank the National Facility for Gene Function in Health and Disease (supported by a grant from the Department of Biotechnology, Government of India; BT/INF/<u>22/SP17358/2016</u>) at IISER Pune for maintaining and providing mice and zebrafish for this study.

## Ethical approval

The animal studies were carried out in compliance with the guidelines of the Institutional Animal Ethics Committee at IISER Pune and Shiv Nadar Institution of Eminence and were approved by the committees.

## Author contributions

SG and AS designed the research work and wrote the manuscript with intellectual inputs form SJP, GJD and SM. AS performed all the experiments in mammalian cells and prepared libraries for sequencing in mammalian cells, mice and zebrafish. GJD performed staining and quantification for cell viability and cell cycle analysis for all the conditions. AS, GJD and SJP performed analysis of all ChIPseq and RNAseq datasets. AS and GJD analysed the colorectal cancer patient transcriptome datasets. GJD and SM assisted in the mice xenograft assay experiment and sample harvesting. SM performed western blotting and analysis by densitometry, standardization of drug treatments in cell lines and all the functional assays in zebrafish, including imaging, evaluation and quantification of phenotype; and qRT-PCR analysis upon overexpression of *sp1* and *ctnnb1*. KS contributed towards generation of the *apc* mutant line.

## Competing Interests

The authors declare no conflict of interest.

## Supporting information

Sharma et al_supplmentary files

## Acknowledgments

AS was supported by the Senior Research Fellowship from the University Grants Commission, India. GJD was the recipient of a Senior Research Fellowship from the Indian Institute of Science Education and Research (IISER), Pune, India. SJP was supported by the Senior Research Fellowship from the Council of Scientific and Industrial Research, India. SM was supported by the DST-INSPIRE Fellowship for Higher Education, India. The authors thank Dr Soumen Khan for helping with the initial sequencing analysis. The authors thank IISER Pune core facilities for their important contributions to this work, especially the imaging Facility for confocal microscopy. We thank the staff members at the National Facility for Gene Function in Health and Disease (NFGFHD) at IISER Pune and Center for Integrative and Translational Research (CITRES) for maintaining and providing mice and zebrafish for this study. We thank Dr Anil Challa for help in design of the genome editing strategy. Figures 1A, 1E, 6A, 7A, 7G, 8 and S8D were created using BioRender.

## Notes

### Competing Interest Statement

The authors have declared no competing interest.

