## Supplementary material for "SP1 amplifies the effects of cellular proliferation via the Wnt/β-catenin pathway in colon development and cancer": Sharma et al_supplmentary files

### Sharma et al, Supplementary Figure S1

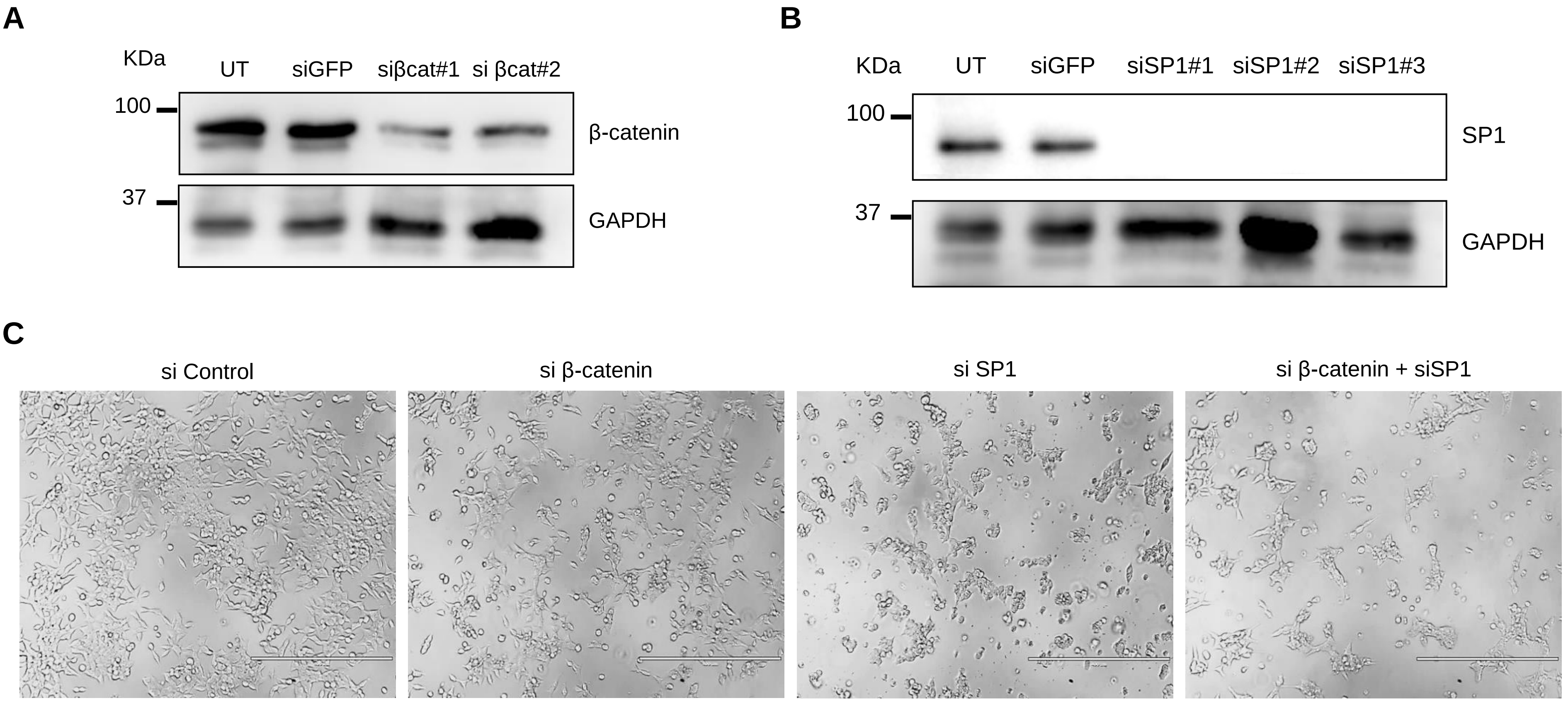


**Supplementary Figure S1: Knockdown efficiency of siRNAs used in the study. A.** Immunoblot analysis to monitor the efficiency of siRNAs against β-catenin. **B.** Immunoblot analysis to assess the efficiency of siRNAs against SP1. **C.** DIC images for cells treated with control siRNA, siβ-catenin, siSP1 and combination of both.

### Sharma et al, Supplementary Figure S2

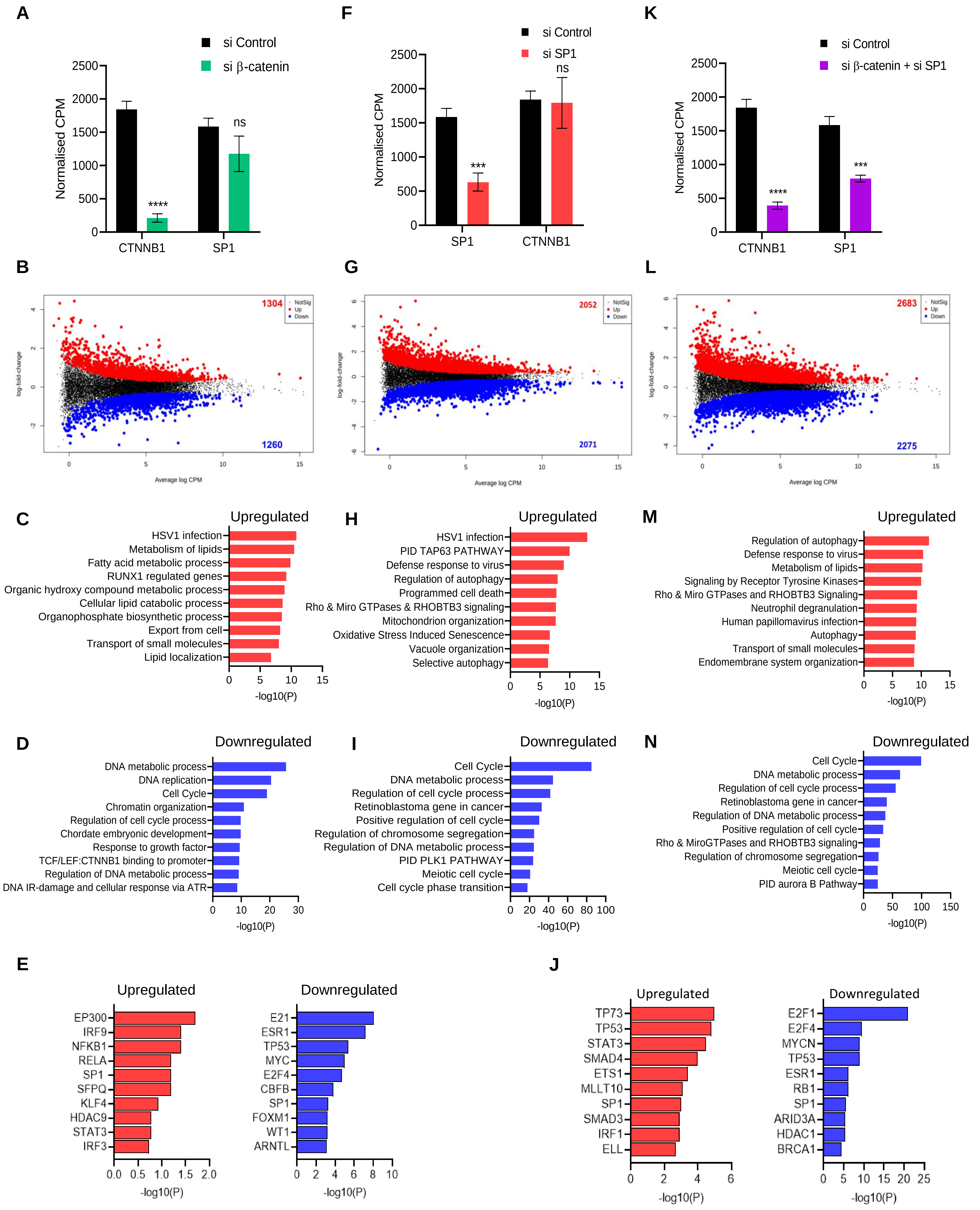

Sharma et al, Supplementary Figure S2

**O**

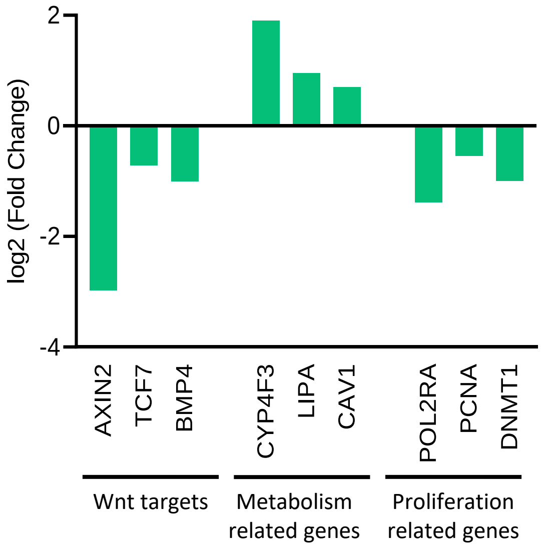


**P**

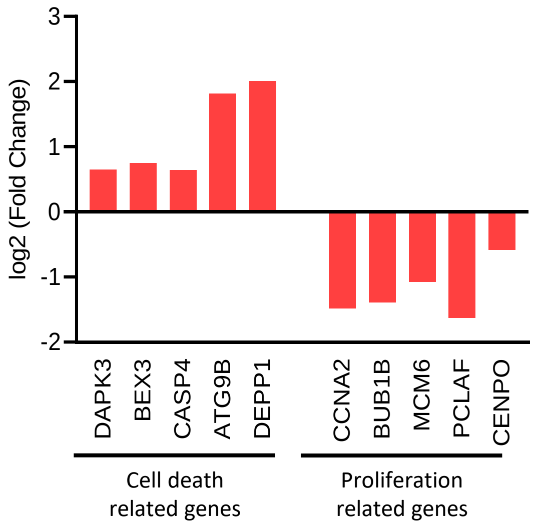


**Supplementary Figure S2: Individual β-catenin and SP1 perturbation dysregulate the expression of genes required for cell growth and survival.** (Upregulated genes are marked in red, and downregulated genes are marked in blue; unpaired t-test was performed to calculate significance for all). **A.** Expression of β-catenin and SP1 mRNA in control and β-catenin siRNA-treated cells. **B.** Volcano plot for the DE genes in β-catenin siRNA-treated cells. **C.** GO analysis for upregulated genes in β-catenin siRNA-treated cells. **D.** GO analysis for downregulated genes in β-catenin siRNA-treated cells. **E.** TRRUST analysis for upregulated genes (red) and downregulated genes (blue) in β-catenin siRNA-treated cells. **F.** Expression of β-catenin and SP1 mRNA in control and SP1 siRNA-treated cells. **G.** Volcano plot for the differentially expressed genes. **H.** GO analysis for upregulated genes in SP1 siRNA-treated cells. **I.** GO analysis for downregulated genes in SP1 siRNA-treated cells. **J.** TRRUST analysis for upregulated genes (red) and downregulated genes (blue) in SP1 siRNA-treated cells. **K.** Expression of β-catenin and SP1 mRNA in control and β-catenin + SP1 siRNA-treated cells (Unpaired t-test was performed to calculate significance). **L.** Volcano plot for the differentially expressed genes in DKD. **M.** Volcano plot for the differentially expressed genes in SP1 siRNA-treated cells. **N.** GO analysis for upregulated genes in SP1 siRNA-treated cells. **O.** log2 Fold change for selected targets of β-catenin in β-catenin siRNA-treated cells. **P.** log2 Fold change plotted for selected target genes of SP1 in SP1 siRNA-treated cells.

Sharma et al, Supplementary Figure S3

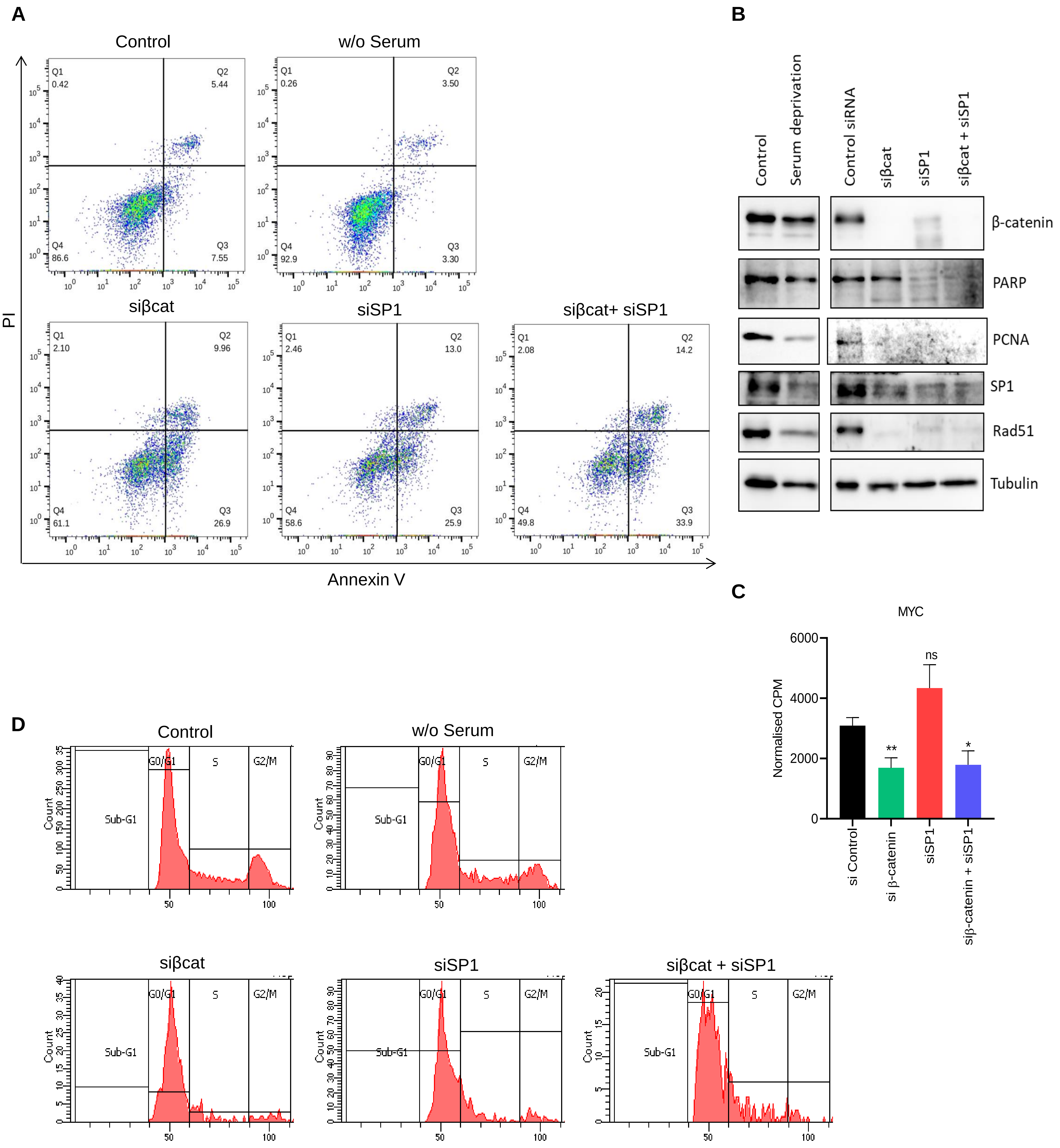


**Supplementary Figure S3: β-catenin and SP1 enhance cell survival and growth. A.** Flow cytometry analysis for cells stained with Annexin V and PI after treatment with control siRNA, siβ-catenin, siSP1 and combination. **B.** WB image for β-catenin, PARP, PCNA, SP1, Rad51 in cells after treatment with control siRNA, siβ-catenin, siSP1, combination and serum deprivation. **C.** Expression of MYC mRNA in control, β-catenin, SP1 and β-catenin + SP1 siRNA treated cells (Unpaired t-test was performed to calculate significance). **D.** Flow cytometry analysis after PI staining for cells treated with control siRNA, siβ-catenin, siSP1 combination and serum deprivation.

Sharma et al, Supplementary Figure S4

**A**

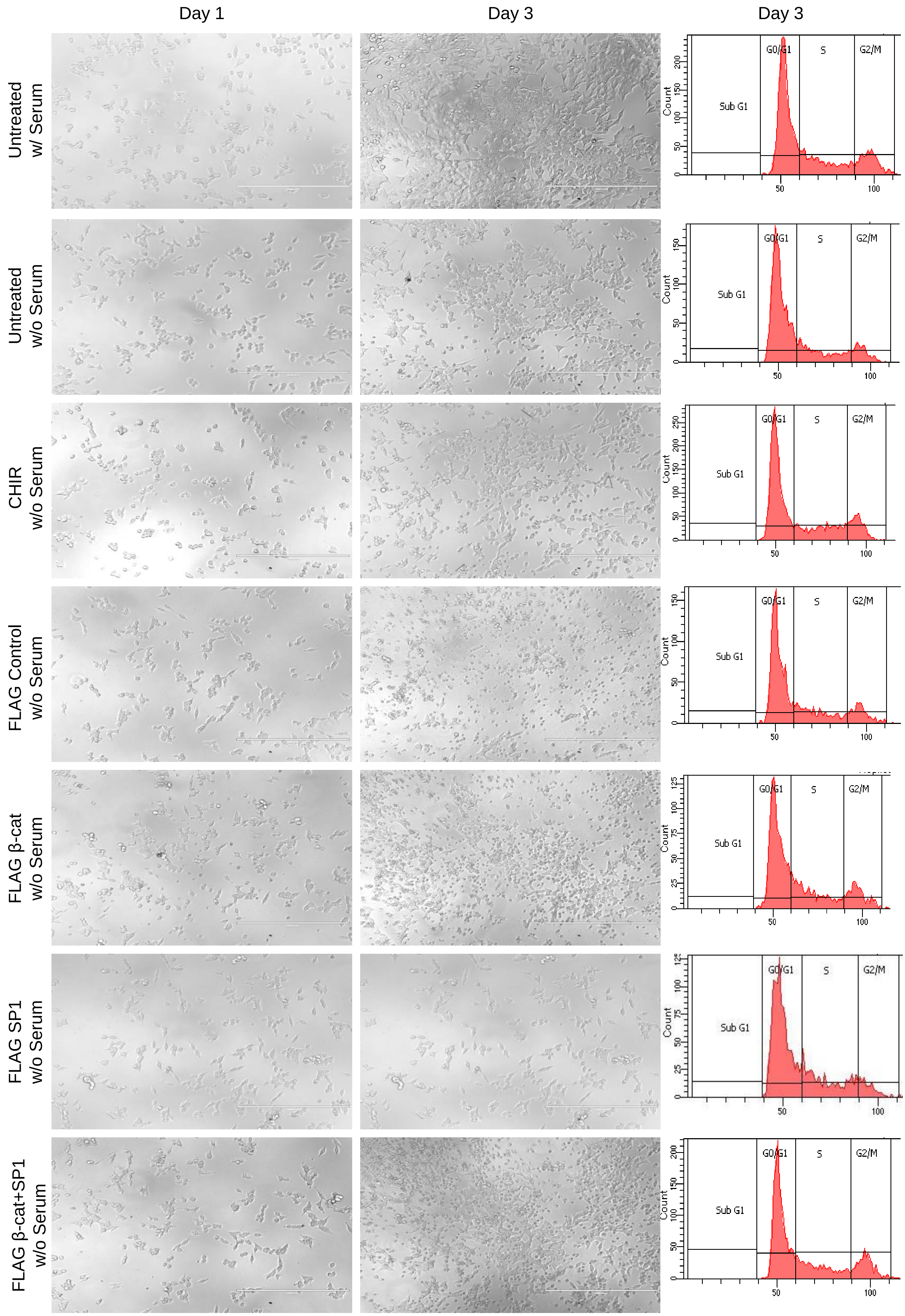

**B**

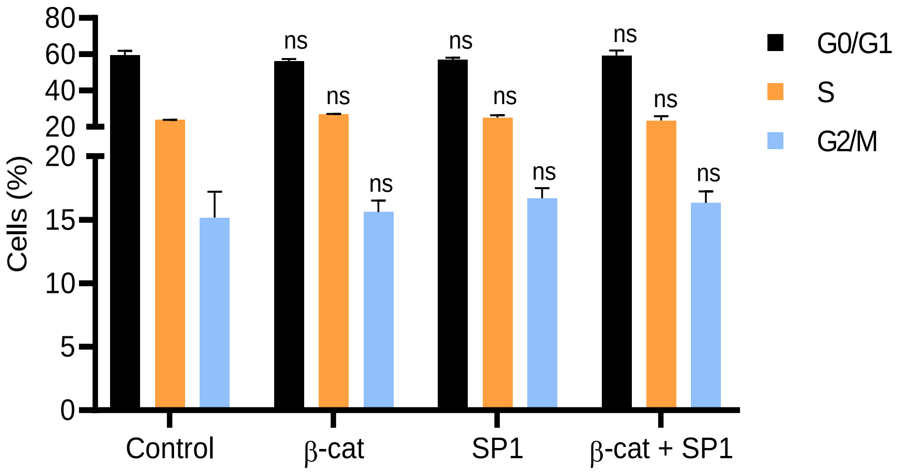


**C**

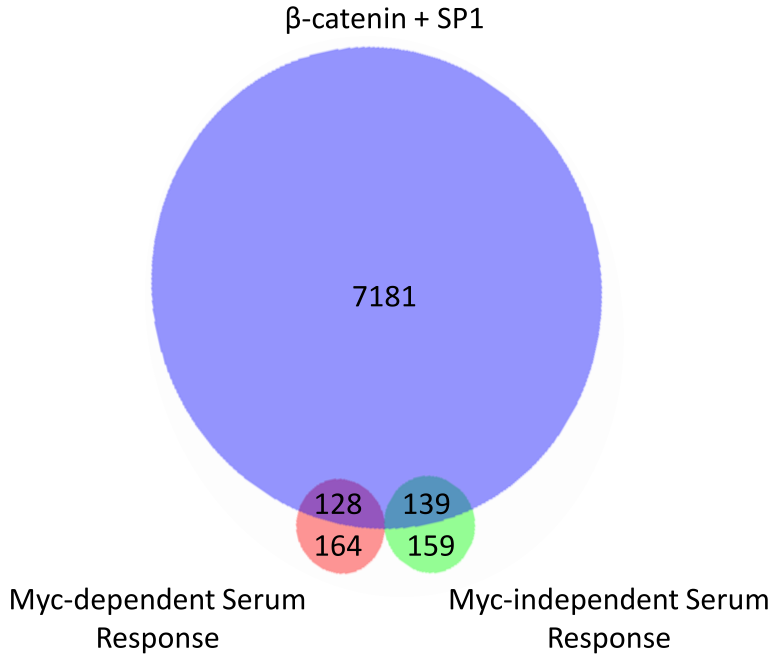


**Supplementary Figure S4: β-catenin and SP1 drive cell growth. A.** DIC images and flow cytometry analysis after PI staining for cells in control medium, serum deprivation, CHIR treatment, overexpression of FLAG control, β-catenin, SP1 and β-catenin + SP1 in serum-deprived medium. **B.** Cell-cycle analysis of cells upon overexpression of FLAG control, β-catenin, SP1 and β-catenin + SP1 in serum-deprived medium. **C.** Overlap of all differentially regulated genes in DKD with Myc-dependent serum-responsive factors and Myc-independent serum-responsive factors.

### Sharma et al, Supplementary Figure S5

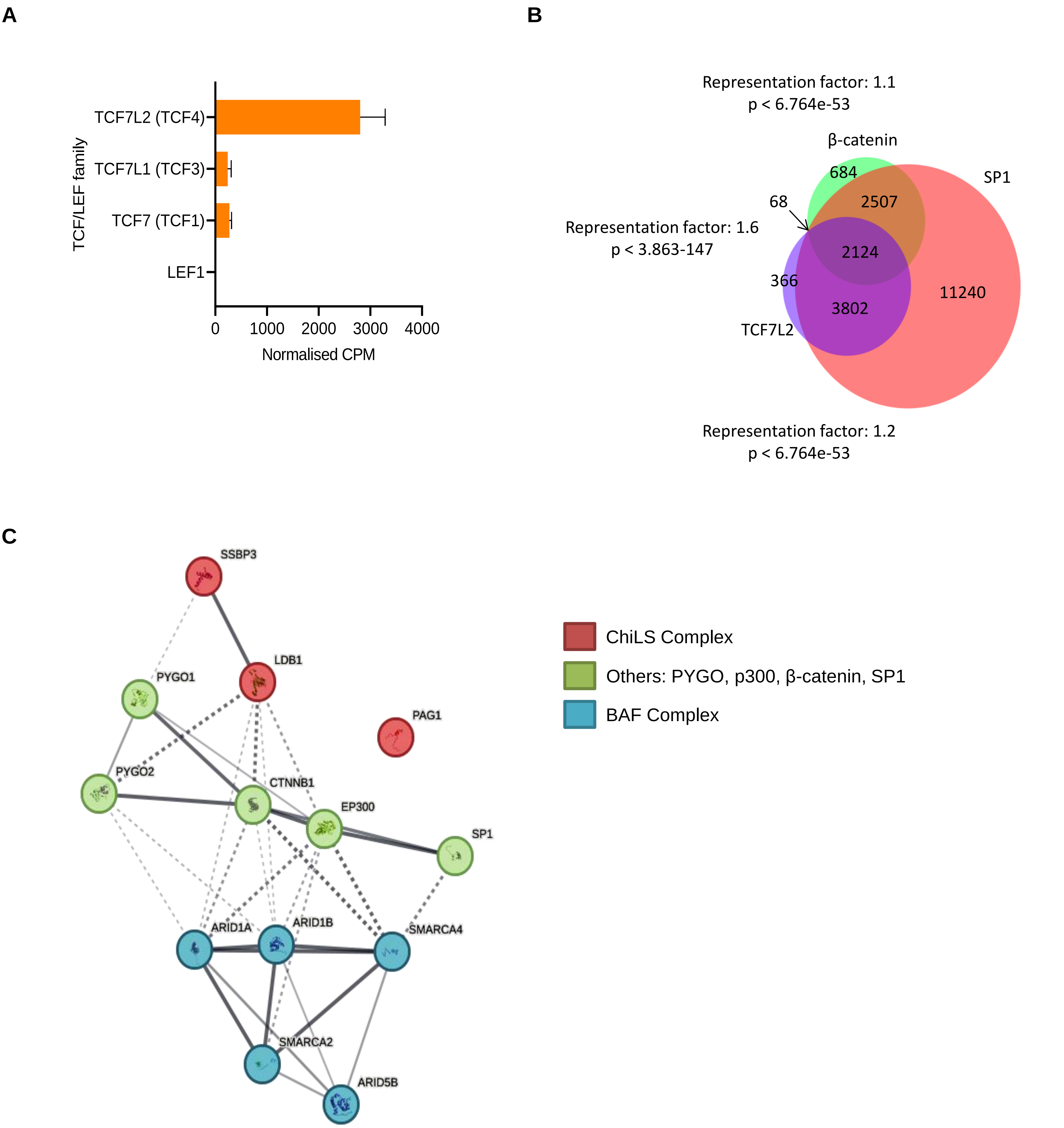


**Supplementary Figure S5: β-catenin and SP1 show chromatin co-occupancy. A.** Expression of all TCF/LEF members in HCT116 cells in control conditions. **B.** Venn diagram to show overlap between the ChIPseq peaks for β-catenin, SP1 and TCF7L2. **C.** STRING network for SP1, β-catenin and the Wnt enhanceosome.

### Sharma et al, Supplementary Figure S6

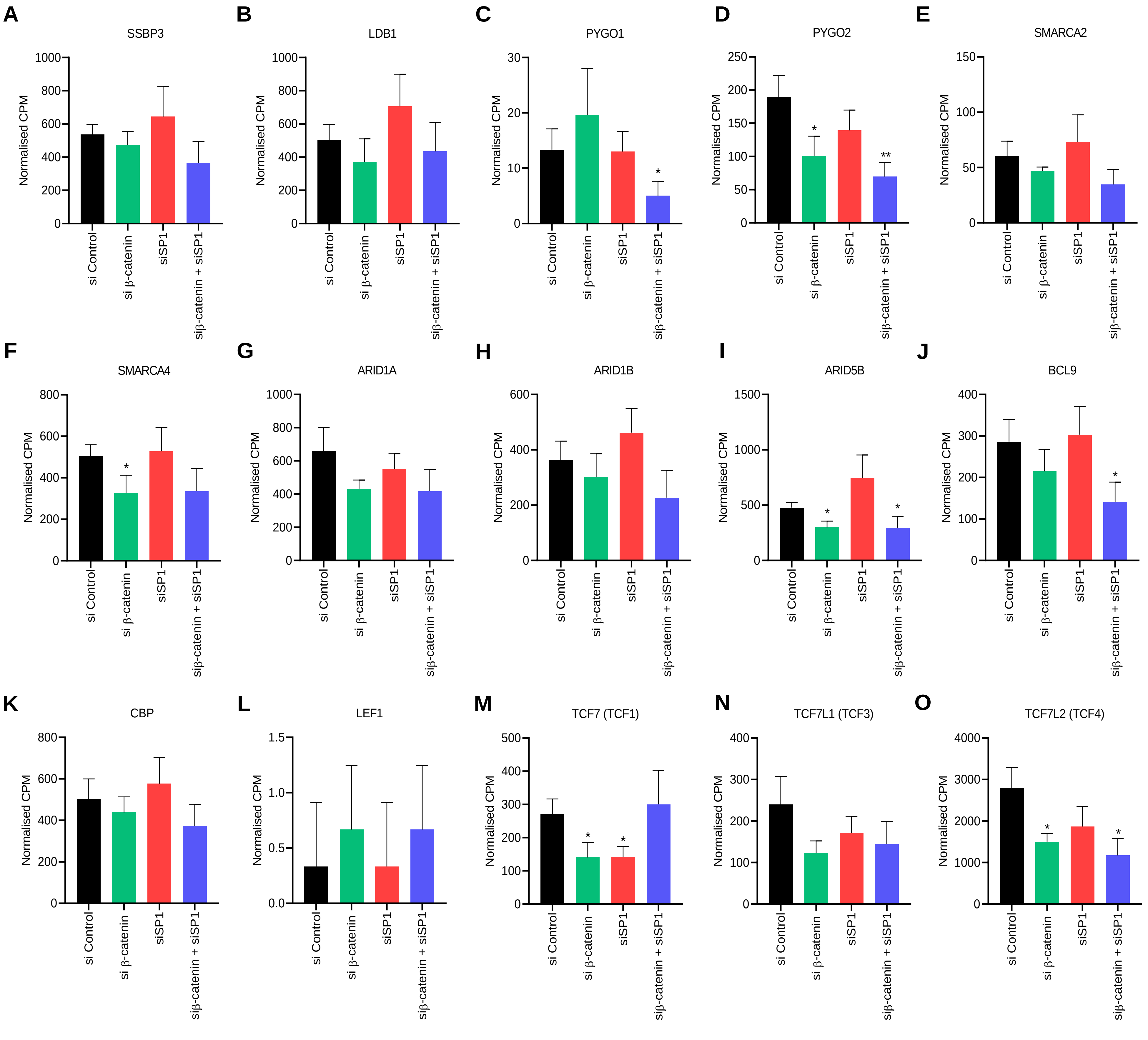


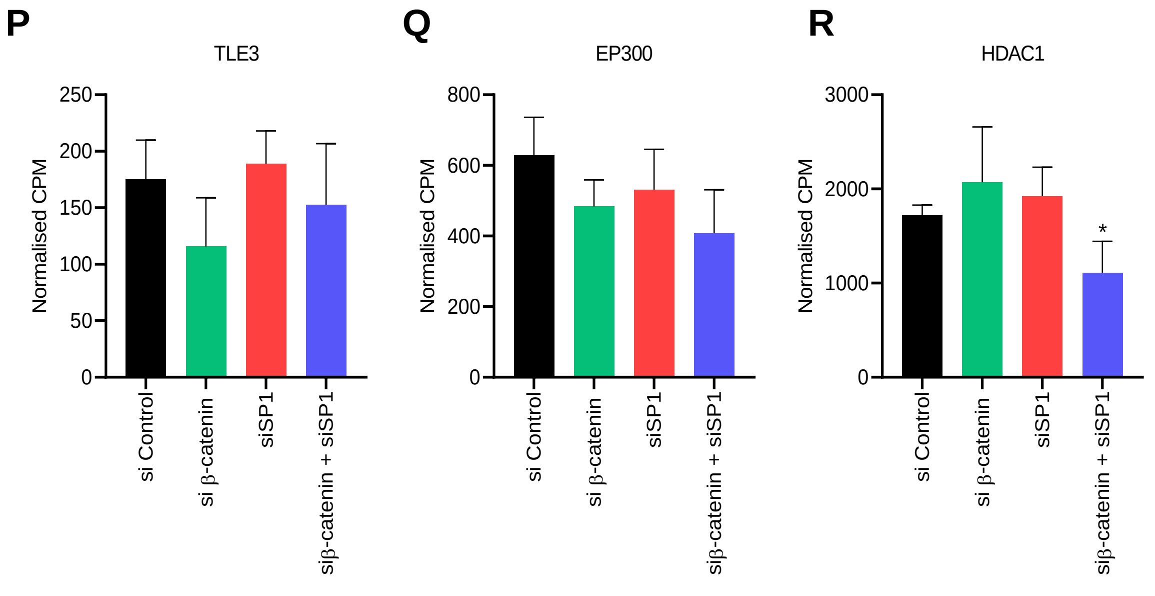


**Supplementary Figure S6: β-catenin and SP1 regulate the expression of the components of the Wnt enhanceosome. A-R:** Expression of mRNA (name mentioned on top for each) in control, β-catenin, SP1 and β-catenin + SP1 siRNA treated cells (Unpaired t-test was performed to calculate significance, results not significant unless mentioned).

### Sharma et al, Supplementary Figure S7

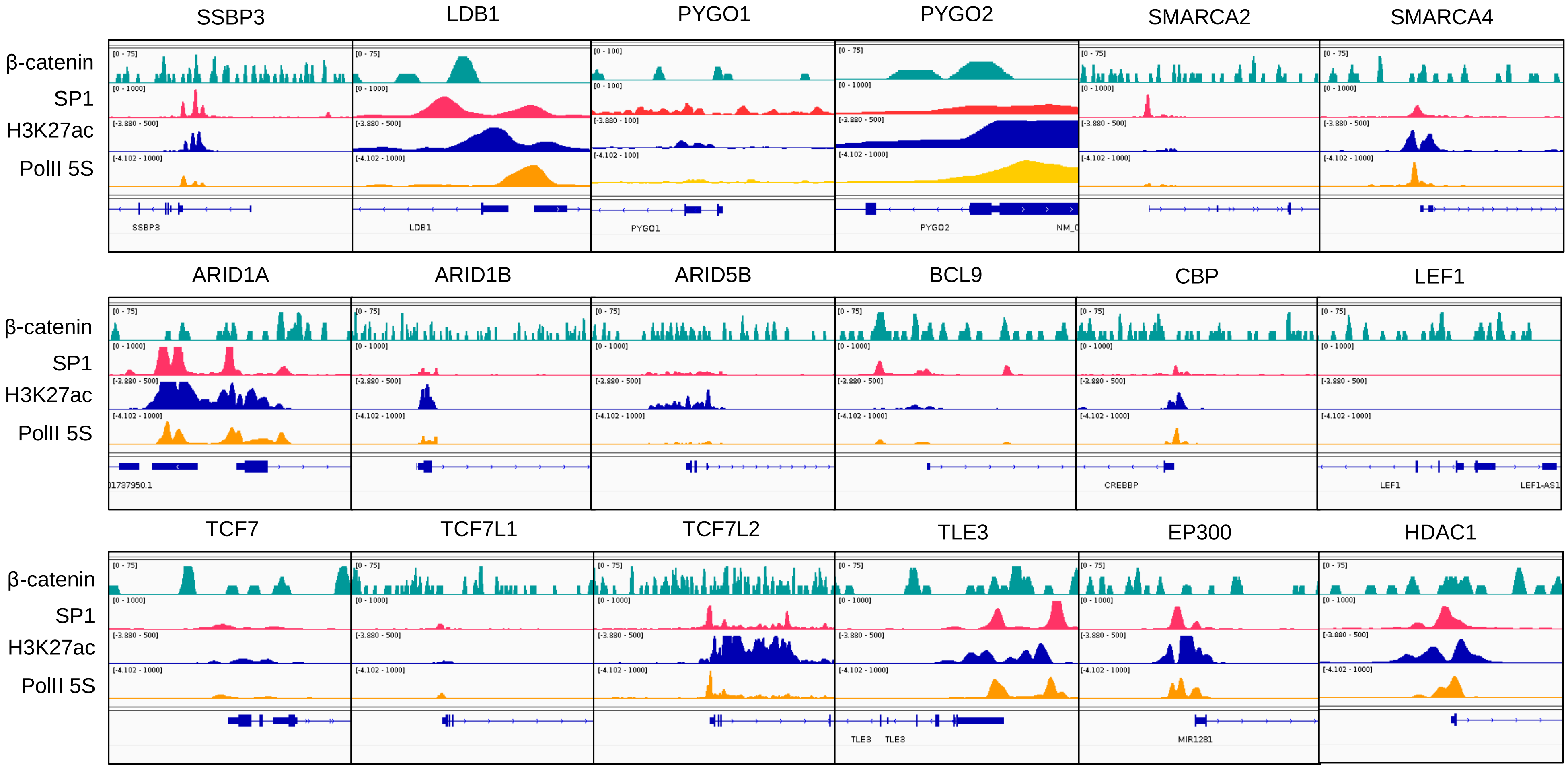


**Supplementary Figure S7: β-catenin and SP1 co-occupy the regulatory regions of the components of the Wnt enhanceosome.** IGV tracks to show occupancy for β-catenin and SP1 on active promoter regions (enrichment for H3K27ac and PolII 5S) of the genes (name mentioned above each panel) for Wnt enhanceosome components.

### Sharma et al, Supplementary Figure S8

A

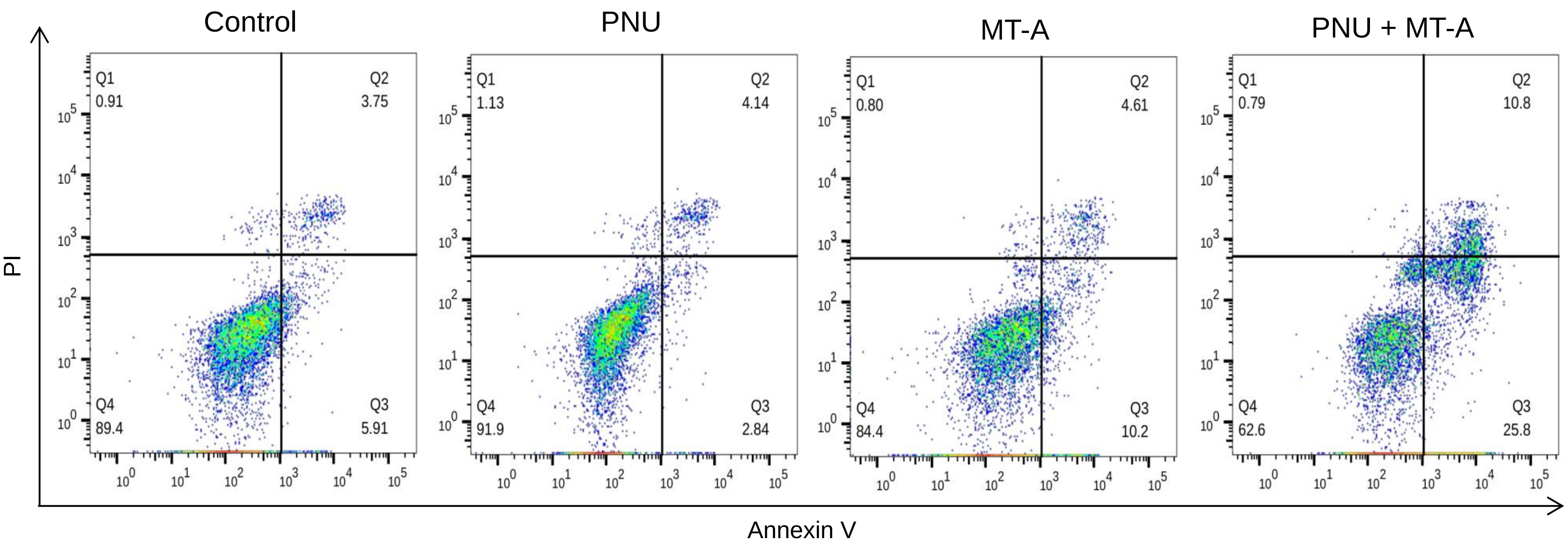


B

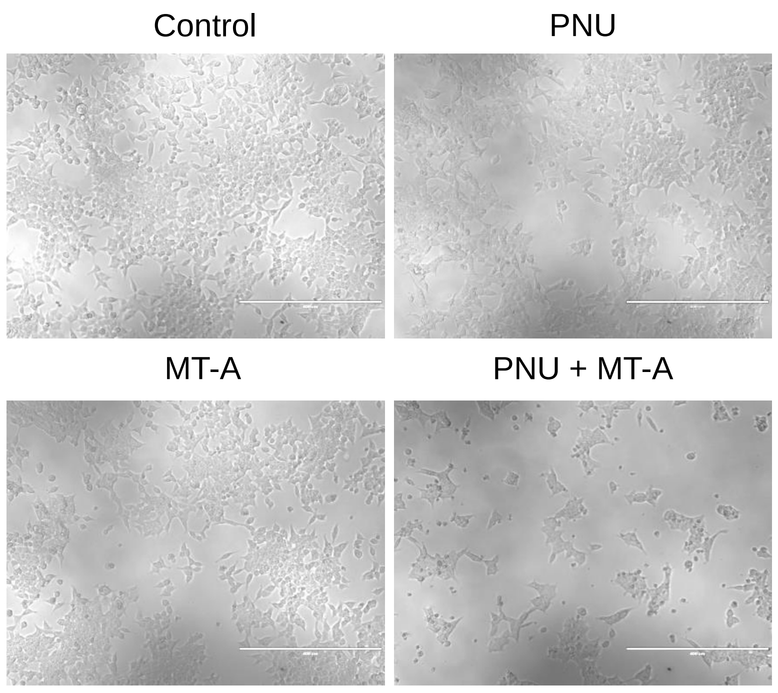


C

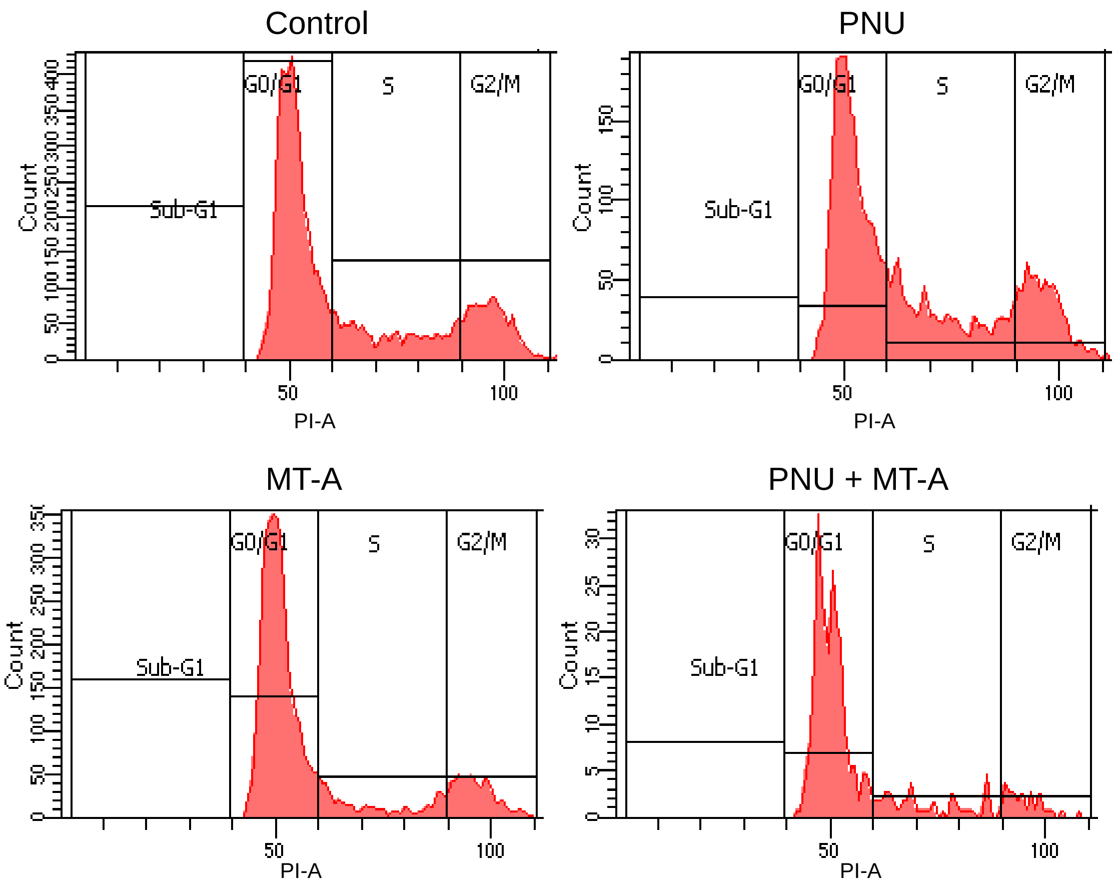


D

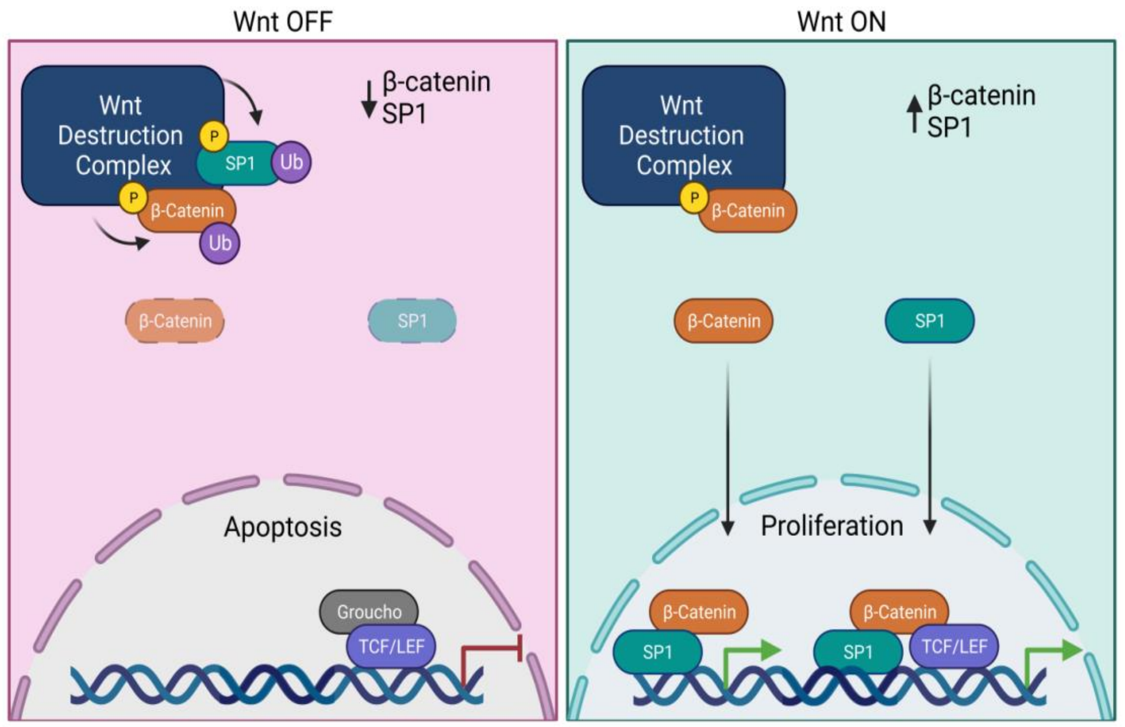


**Supplementary Figure S8: Combinatorial inhibition of β-catenin and SP1 promotes cell death. A.** Flow cytometry analysis of HCT116 cells stained with Annexin V and PI after treatment with PNU, MT-A and PNU + MT-A. **B.** DIC images for cells after treatment with PNU, MT-A and PNU + MT-A. **C.** Flow cytometry analysis after PI staining for HCT116 cells treated with PNU, MT-A and PNU + MT-A. **D.** Model for co-operation between β-catenin and SP1. When the Wnt pathway is off, the Wnt destruction complex primes SP1 and β-catenin for degradation, which in turn causes apoptosis. When Wnt signaling gets activated, the stabilised SP1 and β-catenin translocate into the nucleus. Further, SP1 and β-catenin co-occupy active promoter and enhancer regions with and without TCF, respectively, and activate the expression of genes required for cell proliferation.

### Sharma et al, Supplementary Table 1

| Reagent | Source | Sequence/Identifier |
|---|---|---|
| **siRNAs** | | |
| Control siRNA | Mir et al., 2018 | AACGTACGCGGAATACTTC |
| β-catenin siRNA #1 | Azzolin et al., 2012 | GTAGCTGATATTGATGGAC |
| β-catenin siRNA #2 | Azzolin et al., 2012 | GCTTGGAATGAGACTGCTG |
| SP1 siRNA #1 | Dharmacon | GCCAATAGCTACTCAACTA |
| SP1 siRNA #2 | Dharmacon | GAAGGGAGGCCCAGGTGTA |
| SP1 siRNA #3 | Dharmacon | GGGCAGACCTTTACAACTC |
| **Morpholinos** | | |
| Control | GeneTools | NA |
| sp1 | GeneTools | AAACAGTACCCCCGGACTGACCTGG |
| **IVT primers** | | |
| sp1 | This study | (F)GAGTAATACGACTCACTATAGATGTCAGACTTGAAGAAGCGTG<br>(R) CTAGAATCCGTTGCTGCCG |
| **Public Datasets (in HCT116)** | | |
| ChIPseq Input for SP1 | ENCODE Consortium | ENCSR000BVT |
| ChIPseq IP for SP1 | ENCODE Consortium | ENCSR000BSF |
| ChIPseq Input for TCF4 | ENCODE Consortium | ENCSR000EUX |
| ChIPseq IP for TCF4 | ENCODE Consortium | ENCSR000EUV |
| ChIPseq Input for TEAD4 | ENCODE Consortium | ENCSR000BVT |
| ChIPseq IP for TEAD4 | ENCODE Consortium | ENCSR000BVJ |
| ChIPseq Input (HMs) | ENCODE Consortium | ENCSR000BMK |
| ChIPseq IP for H3K4me1 | ENCODE Consortium | ENCSR161MXP |
| ChIPseq IP for H3K4me3 | ENCODE Consortium | ENCSR333OPW |
| ChIPseq IP for H3K27ac | ENCODE Consortium | ENCSR661KMA |
| ChIPseq Input for POL25S | ENCODE Consortium | ENCSR000BMK |
| ChIPseq IP for POL25S | ENCODE Consortium | ENCBS389ENC |
| ChIPseq Input for β-catenin | Bottomly et al., 2010 | SRA012054 |
| ChIPseq Input for BRG1 | Mathur et al., 2017 | SRR2133624 |
| ChIPseq IP for BRG1 | Mathur et al., 2017 | SRR2133616 |
| ChIPseq for LDB1 | Lai et al., 2020 | SRR11969104 |
| ChIPseq Input for TEAD1 | Lai et al., 2020 | SRR6476373 |
| ChIPseq IP for TEAD1 | Lai et al., 2020 | SRR6476317 |

### Sharma et al, Supplementary Table 2

| Reagent | Source | Identifier |
|---|---|---|
| **Experimental models** | | |
| Cell line HCT116 | ECACC | 91091005 |
| Cell line HEK293T | ECACC | 12022001 |
| Mice NOD.SCID | Envigo Research | NOD.CB17-Prkdc$^{scid}$/NCrHsd |
| Zebrafish WT AB | Heisenberg lab | NA |
| Zebrafish WT Tübingen (TU) | Sonawane lab | NA |
| Zebrafish apc$^{-/-}$ isolated RNA (5dpf) | Kirankumar lab | NA |
| Zebrafish WT sibling isolated RNA (5dpf) | Kirankumar lab | NA |
| **Recombinant Constructs** | | |
| pCMV10 FLAG-SP1 WT | Mir et al., 2018 | NA |
| pCMV9 FLAG-SP1 MT | Mir et al., 2018 | NA |
| pCMV10 FLAG-β-catenin WT | Mir et al., 2018 | NA |
| pSC2+ ctnnb1 | Heisenberg lab | NA |
| **Antibodies** | | |
| SP1 | Cell Signaling Technology | 9389 |
| β-catenin | BD Transduction Labs | 610153 |
| YAP | Santa Cruz | sc-101199 |
| Tubulin | BIO-RAD | MCA4403Z |
| GAPDH | Santa Cruz | |
| BRG1 | Santa Cruz | sc-10768 |
| TCF7L2 | Cell Signaling Technology | 2569 |
| p300 | Santa Cruz | sc-48343 |
| PARP | Cell Signaling Technology | 9532 |
| PCNA | Santa Cruz | sc-25280 |
| Rad51 | Santa Cruz | sc-8349 |
| Normal Rabbit IgG | Cell Signaling Technology | 2729 |
| Normal Mouse IgG | Upstate | 12-371 |
| **Proteins and chemicals** | | |
| Human Wnt3A ligand | R&D Systems | 5036-WN |
| Human Fibronectin | Merck | F2006 |
| DAPI | Merck | D9542 |
| PNU-74654 | Cayman Chemical | Cay16349-10 |
| Mithramycin-A | Cayman Chemical | Cay11434-5 |
| CHIR | Calbiochem | 531167 |
